# SILAC-based quantitative proteomics reveals pleiotropic, phenotypic modulation in primary murine macrophages infected with the protozoan pathogen *Leishmania donovani*

**DOI:** 10.1101/742841

**Authors:** Despina Smirlis, Florent Dingli, Pascale Pescher, Eric Prina, Damarys Loew, Najma Rachidi, Gerald F. Späth

**Affiliations:** Unité de Parasitologie Moléculaire et Signalisation, Institut Pasteur and Institut National de Santé et Recherche Médicale INSERM U1201, Paris, France; Molecular Parasitology Laboratory, Hellenic Pasteur Institute, Athens, Greece; Laboratoire de Spectrométrie de Masse Protéomique, Centre de Recherche, Institut Curie, Université de recherche PSL, Paris, France

**Keywords:** quantitative proteomics, bone marrow-derived macrophages, *Leishmania donovani*, SILAC, LC-MS/MS

## Abstract

Leishmaniases are major vector-borne tropical diseases responsible for great human morbidity and mortality, caused by protozoan, trypanosomatid parasites of the genus *Leishmania.* In the mammalian host parasites survive and multiply within mononuclear phagocytes, especially macrophages. However, the underlying mechanisms by which *Leishmania* spp affect their host, are not fully understood. Herein, proteomic alterations of primary bone marrow-derived, BALB/c macrophages are documented after 72 h of infection with *Leishmania donovani* insect-stage promastigotes, with the use of a SILAC-based, quantitative proteomics approach. The protocol was optimised by combining strong anion exchange and gel electrophoresis fractionation that displayed similar depth of analysis (>5500 proteins). Our analyses revealed 86 differentially modulated proteins (35 showing increased and 51 decreased abundance) in response to *Leishmania donovani* infection. The proteomics results were validated by analysing the abundance of selected proteins. Intracellular *Leishmania donovani* infection led to changes in various host cell biological processes, including primary metabolism and catabolic process, with a significant enrichment in lysosomal organisation. Overall, our analysis allows new technical insight into the challenges of quantitative proteomics applied on primary cells, and establishes the first proteome of *bona fide* primary macrophages infected *ex vivo* with *Leishmania donovani*, revealing new mechanisms acting at the host/pathogen interface.

## INTRODUCTION

The leishmaniases include a group of diseases caused by more than 21 protozoan species of the trypanosomatid genus *Leishmania*, which are transmitted by the bite of phlebotomine sandflies [1]. These diseases, prevalent in 98 countries, have different clinical outcomes that range from localised skin ulcers (cutaneous leishmaniasis) to lethal systemic disease (visceral leishmaniasis, VL)[2]. VL is the most serious form of the disease, with 200 000 to 400 000 new cases documented every year [2] and a lethal outcome if left untreated. VL is caused mainly by strains of the *Leishmania (L.). donovani* complex and is characterised by systemic symptoms, such as fever and weight loss [3].

All *Leishmania* parasites share a digenetic life cycle and live either as extracellular, motile promastigotes within the midgut of a sand fly vector, or as intracellular, non-motile amastigotes in professional phagocytes, with macrophage phagolysosomes representing the primary niche for proliferation [4]. These phagocytic mononuclear cells carry important, immuno-regulatory functions and accordingly can differentiate into phenotypically distinct subtypes [5], including pro-inflammatory M1 cells with microbicidal properties and alternatively activated, anti-inflammatory M2 cells involved in tissue repair (reviewed in [6]). These M1 and M2 phenotypes represent the extremes of a continuous spectrum of polarisation states [7]. Even though both the initial *Leishmania* infection through the bite of infected sand flies and the later acute stages of the disease are characterised by inflammation of the infected tissue [8, 9], *Leishmania* thrives in these environments by efficiently dampening the pro-inflammatory response of its infected host cell, possibly by exploiting developmental programs underlying macrophage phenotypic plasticity [10].

The prevention of *Leishmania* clearance and persistent infection of macrophages is governed by the expression of specific parasite virulence factors, such as highly abundant parasite surface glycolipids [lipophosphoglycan (LPG) and glycoinositol phospholipids (GIPLs)], or the GPI anchored metalloprotease, GP63 [11–15]. These and other factors have been shown to remodel the macrophage phenotype to establish permissive conditions for *Leishmania* intracellular survival. Initially, following entry into the macrophage, *L. donovani* promastigotes delay phagosomal/lysosomal fusion and phagosome maturation [16–18], and subsequently modify host cell signaling and metabolic processes, inhibit apoptosis and suppress antigen presentation (reviewed in [10]).

However, this ideal concept of ‘macrophage deactivation’ upon *Leishmania* infection is clearly an oversimplification, as the macrophage response to infection under clinically more relevant conditions can depend on many parameters, including the host cell differentiation state, the tissue context, and parasite species or even strain [19, 20]. Systems-level, ‘omics’ approaches revealed the complex interface between the parasite and the macrophage and shed important new light on the host cell response to *Leishmania* infection, particularly at the transcript level [8, 9, 19, 21–27]. However, transcript abundance does not always correlate with protein levels and thus may not give an accurate picture of the cellular phenotype. Proteome analyses certainly allows for a more relevant functional insight – but the few proteomics screens performed on *Leishmania* infected cells used either immortalised, macrophage-like cell lines [28] known to be phenotypically and functionally different from *bona fide* macrophages [29–32], or a semi-quantitative approach based on spectral counting [20] that lacks quantitative performance [33]. Here we overcame these limitations and for the first time quantified proteomic changes in primary, bone marrow-derived macrophages (BMDMs) after 72 h of infection with virulent *L. donovani* promastigotes using stable isotope labeling by <u>a</u>mino acids in <u>c</u>ell culture (SILAC). SILAC, a technique that relies on the metabolic replacement of a given natural amino acid (referred to as ‘light’) with a non-radioactive, stable isotope (referred to as ‘heavy’) [34], is considered one of the most accurate quantitative approaches when using cell culture systems to study dynamics of biological processes [33, 34]. Our analysis provides new insight into *L. donovani*-induced phenotypic changes of primary macrophages and mechanisms of intracellular parasite survival acting at the host/pathogen interface.

## MATERIALS AND METHODS

### Ethics statement

All animal experiments complied with the ARRIVE guidelines and were carried out according to the EU Directive 2010/63/EU for animal experiments. All animals were housed in an A3 animal facility of Institute Pasteur. Housing and conditions were in compliance with the protocol approved by the Pasteur Institute ethical committee for animal experimentation (CETEA: C2TEA 89) and procedures were conducted in agreement with the project licenses HA0005 and #10587, issued by the Pasteur Institute CETEA and the Ministère de l’Enseignement Supérieur de la Recherche et de l’Innovation (MESRI) respectively.

### Leishmania culture, BMDM differentiation and infection

*L. donovani* strain 1S2D (MHOM/SD/62/1S-CL2D obtained from Henry Murray (Weill Cornell Medical College, New York, USA) was maintained in female RjHan:AURA golden Syrian hamsters between four and six weeks of age (Janvier Labs) by serial *in vivo* passaging. Amastigotes were purified from hamster spleens as described [35]and differentiated into promastigotes in culture that were maintained for no more than 5 *in vitro* passages. Parasites were cultivated at 26°C in fully supplemented M199 medium (10% FCS, 25□mM HEPES, 4□mM NaHCO_3_, 1□mM glutamine, 1 × RPMI 1640 vitamin mix, 0.2□μM folic acid, 100□μM adenine, 7.6□mM hemin, 8□µM biopterin, 50 U ml^−1^ of penicillin and 50□μg ml^−1^ of streptomycin, pH7.4).

Bone marrow cell suspensions were recovered from tibias and femurs of BALB/c mice (Janvier) and suspended in bacteriologic Petri dishes (Greiner, Germany) at a concentration of 1.35×10^6^ ml^−1^ in complete medium i.e. RPMI 1640 supplemented with 50 μM β-mercaptoethanol, 15% heat-inactivated fetal calf serum (GIBCO), 50 U ml^−1^ of penicillin and 50□μg ml^−1^ of streptomycin and 75 ng mL^−1^ of macrophage Colony Stimulating Factor-1 (mCSF-1) (Immunotools). Cells were incubated at 37°C and 5% CO_2_. Three days after plating, 0.4 volume of fresh complete medium was added to the cells. Six days later, adherent bone marrow-derived macrophages (BMDMs) were washed with Dulbecco’s phosphate buffered solution (PBS). Cells were detached by incubation with Dulbecco’s PBS without Ca^2+^ and Mg^2+^ (Biochrom AG, Berlin, Germany) containing 25 mM EDTA for 30min at 37°C.

For *Leishmania* infection, recovered BMDMs were plated in flat-bottom 6-well plates (Tanner, Switzerland) at a density of 4×10^6^ cells per well and were maintained in complete medium. Six hours after plating, macrophages were infected with 4×10^7^ *L. donovani* stationary phase promastigotes (multiplicity of infection of 10:1, *in vitro* passage 2 after differentiation from splenic amastigotes) for 4 h and incubated at 37°C in a 5% CO_2._ Following infection, macrophages were washed four times with PBS to remove free promastigotes. Infected and non-infected macrophages were then maintained for 72 h in complete medium containing only 25ng mL^−1^ mCSF-1.

### SILAC labelling and sample preparation

For labelling cells by SILAC, equal numbers of BMDMs were differentiated and cultured as described above, but using RPMI 1640 without Lysine and Arginine (Thermo Fisher Scientific) that was supplemented either with natural amino acids (L-Lysine, 0.274 mM; L-Lysine, 1.15 mM; Arginine, 1.15 mM) to obtain “light” cells (referred to as control) or with amino acid isotopes ^2^H_4_-Lysine (Lys4) and ^13^C_6-_-Arginine (Arg6) at the same concentrations to obtain “heavy” cells (referred to as labelled) (Thermo Scientific). Fetal calf serum (GIBCO) was dialysed over night against 100 volumes of sterile PBS, followed by two new rounds of 3 h dialysis, each against a low cutoff membrane (3.5 kD, Gebaflex). The light or heavy RPMI 1640 was supplemented with 15% (v/v) of dialysed and filter-sterilised serum, 50 μM β-mercaptoethanol, 50 U ml^−1^ penicillin, 50□μg ml^−1^ streptomycin, and 75 ng mL^−1^ of mCSF-1.

BMDMs cultivated in SILAC medium were washed three times in PBS and were lysed with a buffer containing 8M urea, 50 mM ammonium bicarbonate pH7.5, one tablet per 10 ml of cOmplete^™^ Protease Inhibitor Cocktail tablets (Sigma), and one tablet of the phosphatase inhibitor cocktail PhosSTOP (Roche). Cell extracts were incubated on ice 30 min, sonicated 5 min, and centrifuged 15 min at 14,000 *g* to eliminate cell debris. Proteins were quantified in the supernatants using the RC DC™ protein assay kit (Bio-Rad), according to the manufacturer’s instructions. For protein identification and quantification, control infected were mixed with labeled non-infected and labeled infected were mixed with control non-infected at an equal ratio.

### SILAC Sample processing

Samples supernatants as processed above were fractionated by two different methods: (i) polyacrylamide gel electrophoresis fractionation (GEL) and (ii) Strong Anion eXchange (SAX). For GEL fractionation, mixed samples were separated by polyacrylamide gel electrophoresis (PAGE) with the use of Invitrogen NuPAGE 10% Bis-Tris 1.0mm*10 Well gels. Proteins were recovered from 9 gel slices following in-gel digestion as described in standard protocols. Briefly, following the SDS-PAGE and washing of the excised gel slices, proteins were reduced by adding 10 mM DTT (Sigma Aldrich) prior to alkylation with 55 mM iodoacetamide (Sigma Aldrich). After washing and dehydrating the gel pieces with 100% acetonitrile, trypsin (Sequencing Grade Modified, Roche Diagnostics) was added and proteins were digested overnight in 25 mM ammonium bicarbonate at 30°C.

SAX-based samples were digested in solution prior to fractionation of peptides. Briefly, the protein extracts obtained by urea extraction were mixed at a 1:1 ratio and subjected to the following process: First, samples were reduced using 10mM DTT, alkylated with 55mM iodoacetamide and digested using Trypsin overnight. Second, peptides were desalted through Sep-Pak C18 cartridges (Waters) and dried down prior to their fractionation. Peptide fractionation was carried out through SAX using in-house prepared microcolumns (anion exchange disks, Empore) and a Britton - Robinson buffer (20mM acetic acid, 20mM phosphoric acid, 20mM boric acid) at six decreasing pH values (11, 8, 6, 5, 4, 3) for the elution of peptides [36].

Samples were loaded onto homemade C18 SepPak-packed StageTips for desalting (principle by stacking one 3M Empore SPE Extraction Disk Octadecyl (C18) and beads from SepPak C18 Cartridge (Waters) into a 200 µL micropipette tip). Peptides were eluted with 40/60 MeCN/H_2_O + 0.1% formic acid and lyophilysed under vacuum. Desalted samples were reconstituted in injection buffer (2% MeCN, 0.3% TFA) for LC-MS/MS analysis.

### LC-MS/MS analysis

Online liquid chromatography (LC) was performed with an RSLCnano system (Ultimate 3000, Thermo Scientific) coupled online to an Orbitrap Fusion Tribrid mass spectrometer (MS, Thermo Scientific). Peptides were trapped on a C18 column (75 μm inner diameter × 2 cm; nanoViper Acclaim PepMap^TM^ 100, Thermo Scientific) with buffer A (2/98 MeCN/H_2_O in 0.1% formic acid) at a flow rate of 4.0 µL/min over 4 min. Separation was performed on a 50 cm × 75 μm C18 column (nanoViper Acclaim PepMap^TM^ RSLC, 2 μm, 100Å, Thermo Scientific) regulated to a temperature of 55°C with a linear gradient of 5% to 25% buffer B (100% MeCN in 0.1% formic acid) at a flow rate of 300 nL/min over 100 min for the SAX and the GEL samples. Full-scan MS was acquired in the Orbitrap analyzer with a resolution set to 120,000, a mass range of m/z 400–1500 and a 4 × 10^5^ ion count target. Tandem MS was performed by isolation at 1.6 Th with the quadrupole, HCD fragmentation with normalised collision energy of 30, and rapid scan MS analysis in the ion trap. The MS^2^ ion count target was set to 2 × 10^4^ and only those precursors with charge state from 2 to 7 were sampled for MS^2^ acquisition. The instrument was run in top speed mode with 3 s cycles.

### MS Data Processing and Protein Identification

Data were searched against the Swiss-Prot *Mus musculus* database (012016) using Sequest^HT^ through Thermo Scientific Proteome Discoverer (v 2.1). The mass tolerances in MS and MS/MS were set to 10 ppm and 0.6 Da, respectively. We set carbamidomethyl cysteine, oxidation of methionine, N-terminal acetylation, heavy ^13^C_6_-Arginine (Arg6) and ^2^H_4_-Lysine (Lys4) as variable modifications. We set specificity of Trypsin digestion and allowed two missed cleavage sites.

The resulting files were further processed by using myProMS (v 3.5) [37]. The Sequest HT target and decoy search result were validated at 1% false discovery rate (FDR) with Percolator. For SILAC-based protein quantification, peptides XICs (Extracted Ion Chromatograms) were retrieved from Thermo Scientific Proteome Discoverer. Global Median Absolute Deviation (MAD) normalisation was applied on the total signal to correct the XICs for each biological replicate. Protein ratios were computed as the geometrical mean of related peptides. To estimate ratio significance, a t-test was performed with the R package limma [38] and the false discovery rate has been controlled using the Benjamini-Hochberg procedure [39].

### RNA isolation and RT qPCR

Total RNA was isolated from macrophages using the Nucleospin RNA isolation kit (MACHEREY-NAGEL GmbH, Germany) according to the manufacturers’ instructions. The integrity of the RNA was assessed using the Agilent 2100 Bioanalyzer system. For RNA reverse transcription to first strand cDNA, 1 μg of total RNA was mixed with random hexamers (Roche Diagnostics) and MMLV-RT reverse transcriptase (Moloney Murine Leukemia Virus, Invitrogen Life Technologies), and the reaction was carried according to the manufacturer’s instructions. qPCR was performed with the QuantiTect SYBR Green kit (Qiagen) according to the manufacturers’ instructions in a 10 μL reaction mixture containing 1 μL cDNA template and 0.5 μM forward/ reverse primers and 5 µl 2x QuantiTect master mix. The reactions were performed in triplicates in white FrameStar® 384 PCR plates (4titude) on a LightCycler® 480 system (Roche Diagnostics, Meylan, France) and gene expression analysis was performed with the use of qBase [40]. The primers used are listed in **Table S1**. Analysis of PCR data and normalisation over the reference genes Yhwaz and RpL19, were performed as previously described [41].

### Western blot analysis

Total protein extracts were separated on 4–12% Bis-Tris NuPAGE^®^ gels (Invitrogen) and blotted onto polyvinylidene difluoride (PVDF) membranes (Pierce). Western blot analyses were performed as previously described [42]. The following primary antibodies were used at 1:1000 dilution: rabbit anti-GSTm (Abcam, ab178684), anti-Asah-1 (ab74469), anti-Rab3IL1 (Abcam, ab98839), anti-Plxna1 (Abcam, ab23391), anti-Hmox-1 (Abcam, ab13243) and anti-Ctsd (Abcam, ab75852) and the monoclonal anti-β actin 13E5 (Cell Signaling Technologies, #4970). The anti-rabbit and anti-mouse secondary antibodies (Pierce) were diluted 1:20000. Blots were developed using SuperSignal™ West Pico (Pierce) according to the manufacturer’s instructions. Western blots were quantified with Image J software.

### Counting of intracellular parasites

Infected and non-infected macrophages plated on coverslips were fixed for 20 minutes at RT with 4% (w/v) parafolmaldehyde (PFA) in phosphate buffered saline (PBS) and incubated for 15 min with PBS containing the nuclear dye Hoechst 33342 (0.5 μg/ml). Excess dye was removed by rinsing coverslips in PBS before mounting on a microscope slide using Mowiol (Sigma-Aldrich). Two different cover slips per condition were analysed with a Leica DMI6000B microscope or an inverted microscope equipped with ApoTome (ZEISS).

### Functional enrichment analysis

GO default annotations were parsed from the available NCBI GO information (http://www.ncbi.nlm.nih.gov/Ftp/), using as background the whole annotation from Uniprot *Mus musculus* reference proteome (UP000000589). The GO enrichment analysis for biological process, molecular function and cell component was performed using the proteins listed in **Table 1** and **2** as input. The multiple testing correction was selected to the Benjamini and Hochberg false discovery rate at a significance level of 0.01. Results were visualised using the Cytoscape 3.5.1 software package [43] and the BinGO plug-in [44].

**Table 1.** Macrophage proteins displaying increased abundance after *L. donovani* infection.

| Description | Uniprot ID | Gene | GEL:L-I/C-NI |  |  | GEL:C-I-NI |  |  |
| --- | --- | --- | --- | --- | --- | --- | --- | --- |
|  |  |  | Ratio 1 | p-value | No. Peptides | Ratio 2 | p-value | No. Peptides |
| Glutamate--cysteine ligase regulatory subunit | O09172 | Gclm | 1.28 | 0.16 | 9 | 1.29 | 0.02 | 10 |
| Complement component C1q receptor | O89103 | Cd93 | 1.35 | 0.19 | 8 | 1.76 | 0.02 | 10 |
| Fatty acid-binding protein, adipocyte | P04117 | Fabp4 | 1.3 | 0.02 | 10 | 1.57 | 0.001 | 8 |
| Urokinase-type plasminogen activator | P06869 | Plau | 6.02 | 0.04 | 5 | 5.41 | 0.003 | 6 |
| Protein disulfide-isomerase A4 | P08003 | Pdia4 | 1.35 | 0.01 | 23 | 1.27 | 0.12 | 26 |
| Ferritin heavy chain | P09528 | Fth1 | 1.37 | 0.03 | 23 | 2.23 | 1.47E-07 | 20 |
| Glutathione S-transferase Mu 1 | P10649 | Gstm1 | 1.36 | 0.003 | 26 | 2.53 | 8.21E-07 | 22 |
| Integrin alpha-5 | P11688 | Itga5 | 1.34 | 0.001 | 18 | 1.27 | 0.3 | 18 |
| Protein disulfide-isomerase A3 | P27773 | Pdia3 | 1.39 | 1.60E-08 | 51 | 1.27 | 7.97E-07 | 59 |
| Ferritin light chain 1 | P29391 | Ftl1 | 2.9 | 3.90E-07 | 18 | 2.64 | 0.009 | 5 |
| CD9 antigen | P40240 | Cd9 | 1.41 | 0.16 | 5 | 1.76 | 0.004 | 6 |
| Signal recognition particle receptor subunit beta | P47758 | Srprb | 1.36 | 0.04 | 7 | 1.25 | 0.14 | 8 |
| V-type proton ATPase subunit D | P57746 | Atp6v1d | 1.29 | 0.31 | 9 | 1.37 | 0.003 | 6 |
| 40S ribosomal protein S15a | P62245 | Rps15a | 1.31 | 0.006 | 11 | 1.34 | 0.499 | 7 |
| Histone H4 | P62806 | Hist1h4 | 1.55 | 3.40E-06 | 12 | 1.33 | 0.15 | 14 |
| Plexin-A1 | P70206 | Plxna1 | 1.65 | 3.00E-05 | 35 | 2.46 | 1.81E-10 | 35 |
| Keratin, type I cytoskeletal 9 | Q6RHW0 | Krt9 | ∞ |  | 8 | ∞ |  | 3 |
| Sterol O-acyltransferase 1 | Q61263 | Soat1 | 1.29 | 0.13 | 10 | 1.31 | 0.03 | 13 |
| Transmembrane protein 214 | Q8BM55 | Tmem214 | 1.34 | 0.04 | 5 | 1.87 | 0.21 | 7 |
| Procollagen galactosyl-transferase 1 | Q8K297 | Glt25d1 | 2.53 | 0.12 | 8 | 1.5 | 0.001 | 11 |
| Na<sup>+</sup>/K<sup>+</sup>-transporting ATPase subunit alpha-1 | Q8VDN2 | Atp1a1 | 1.39 | 0.001 | 17 | 1.26 | 0.007 | 22 |
| Ribophorin I | Q91YQ5 | Rpn1 | 1.65 | 0.0003 | 26 | 1.28 | 0.009 | 32 |
| Protein disulfide-isomerase A6 | Q922R8 | Pdia6 | 1.43 | 0.0004 | 19 | 1.31 | 0.05 | 19 |
| N-acylneuraminate cytidyltransferase | Q99KK2 | Cmas | 1.27 | 0.37 | 11 | 1.72 | 0.002 | 10 |
| BolA-like protein 1 | Q9D8S9 | Bola1 | 1.73 | 0.45 | 2 | ∞ |  | 2 |
| Cap-specific mRNA (nucleoside-2'-O)-methyltransferase 1 | Q9DBC3 | Cmtr1 | 1.39 | 0.72 | 2 | ∞ |  | 4 |
| Hypoxia up-regulated protein 1 | Q9JKR6 | Hyou1 | 1.86 | 2.90E-05 | 36 | 1.38 | 0.008 | 45 |
| Succinyl-CoA ligase [ADP-forming] subunit beta, mitochondrial | Q9Z2I9 | Sucla2 | 1.27 | 0.48 | 10 | 1.37 | 0.007 | 17 |
|  |  |  | Ratio 1 | p-value | No. peptides | Ratio 2 | p-value | No. peptides |
| Phytanoyl-CoA dioxygenase, peroxisomal | O35386 | Phyh | ∞ |  | 2 | 1.25 | 0.53 | 2 |
| 26S protease regulatory subunit 6A | O88685 | Psmc3 | 1.34 | 0.56 | 17 | 1.25 | 0.03 | 21 |
| Glutathione S-transferase Mu 1 | P10649 | Gstm1 | 1.41 | 0.14 | 16 | 2.18 | 0.002 | 22 |
| Amine oxidase [flavin-containing] A | Q64133 | Maoa | 1.31 | 0.54 | 15 | 2.24 | 0.005 | 15 |
| Copper-transporting ATPase 1 | Q64430 | Atp7a | 2.61 | 0.63 | 3 | 2.56 | 0.03 | 4 |
| Protein PRRC2A | Q7TSC1 | Prrc2a | ∞ |  | 3 | 3.86 | 0.08 | 3 |
| TBC1 domain family member 22A | Q8R5A6 | Tbc1d2<br>2a | ∞ |  | 2 | 1.89 | 0.48 | 3 |
| Tumor necrosis factor alpha-induced protein 8-like protein 2 | Q9D8Y7 | Tnfaip81<br>2 | ∞ |  | 4 | 1.44 | 0.71 | 2 |

**Table 2.**
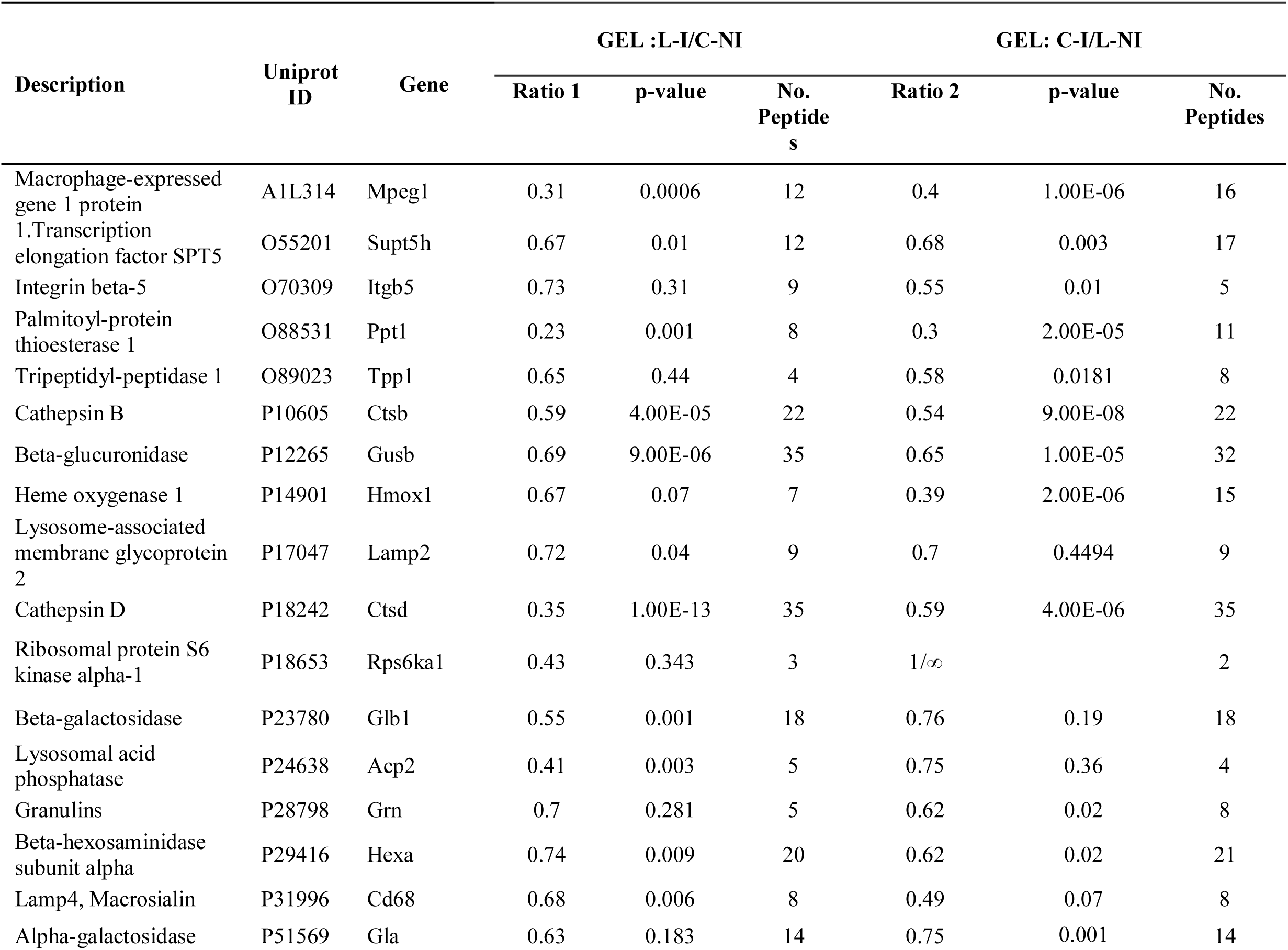
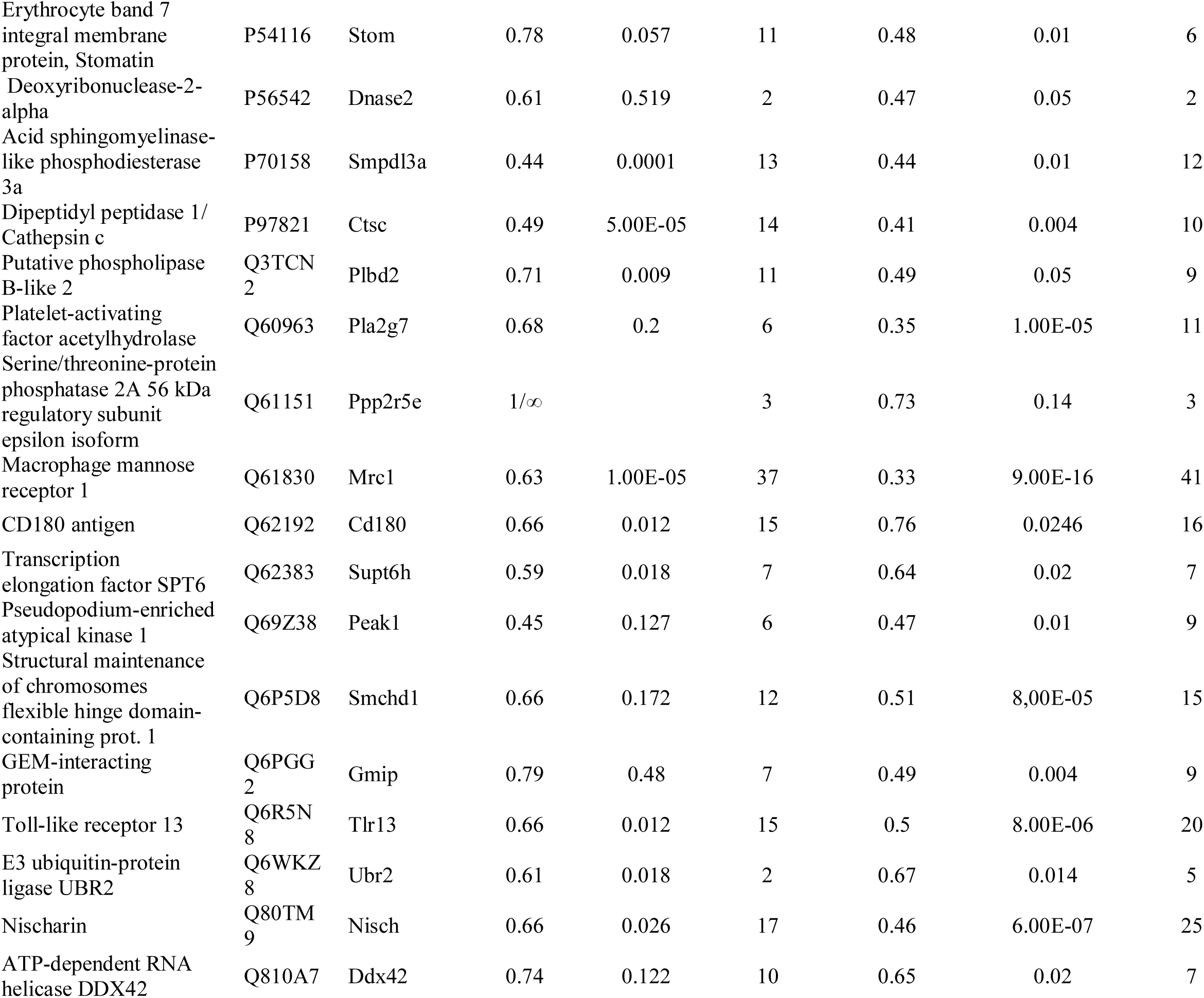
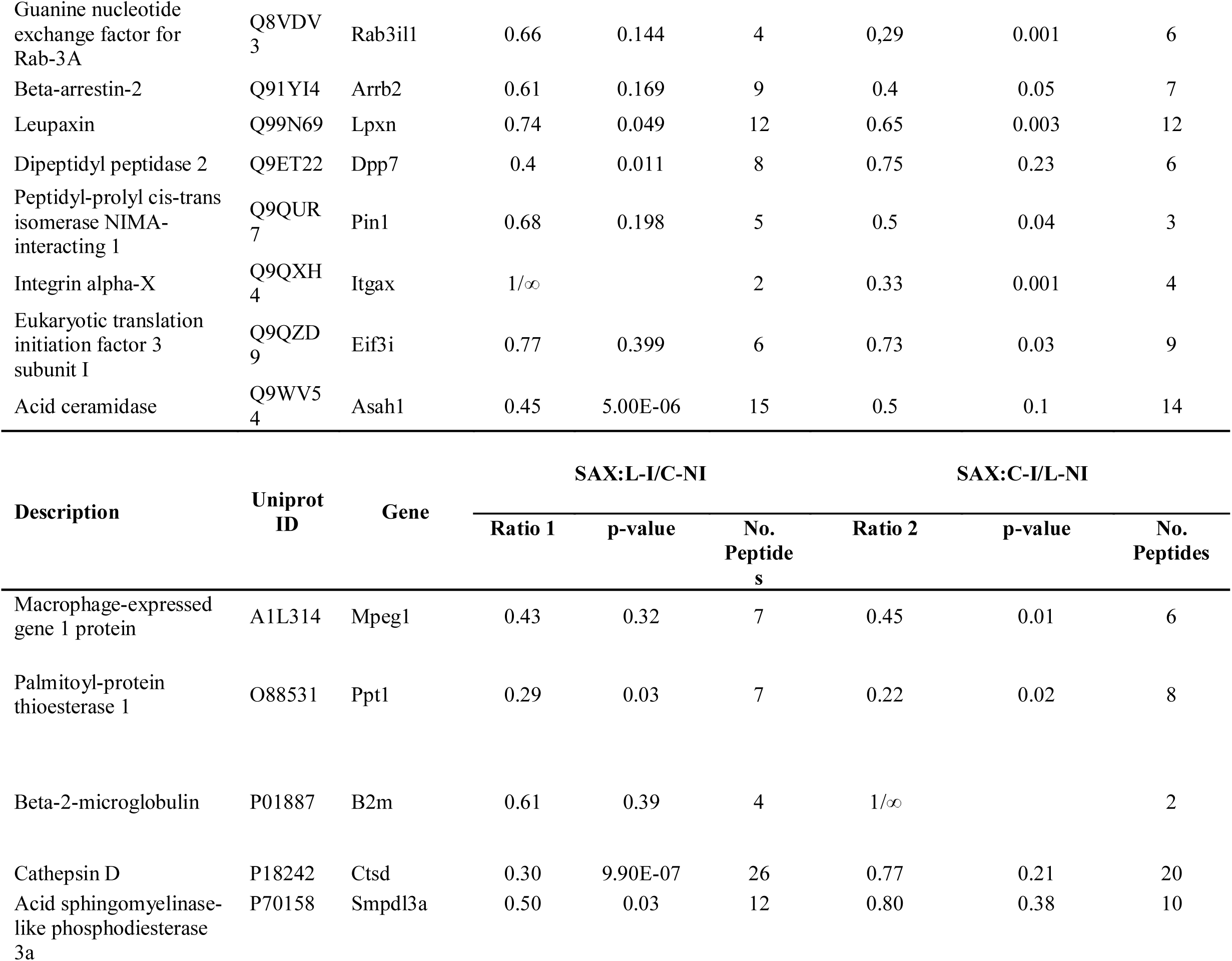
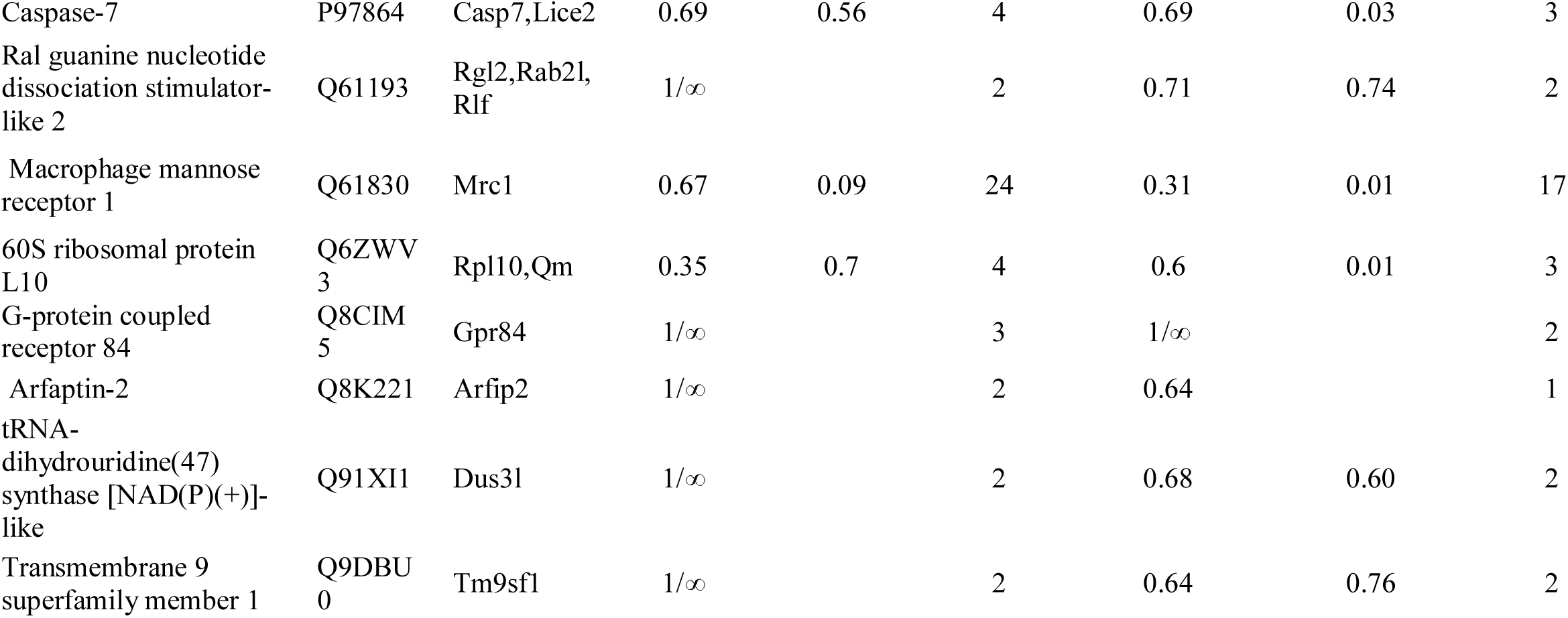
Macrophage proteins displaying decreased abundance after *L. donovani* infection.

## RESULTS AND DISCUSSION

### Metabolic labeling of BMDMs in SILAC medium and experimental workflow for SILAC experiments

First, we tested the capacity of *L. donovani* promastigotes to infect and proliferate within control and isotope-labelled BMDMs. Macrophages at day 6 post-differentiation in SILAC culture medium were infected at a parasite-to-macrophage ratio of 10:1 with *L. donovani* promastigotes at *in vitro* passage 2 after differentiation from lesion-derived amastigotes. Intracellular parasite burden was microscopically assessed at 4 h, 48 h and 72 h post-infection (PI) (**Fig. 1A**). We observed a mean number of 8 parasites per macrophage at 4 h PI, which transiently decreased at 48 h but then recovered at 72 h. More than 80% of macrophages were infected at all time points tested (4 h, 48 h and 72 h, data not shown), indicating robust infection and intracellular parasite growth. Infection was similar in both labelled and control macrophages, suggesting that the metabolic labelling did not influence the ability of parasites to survive and proliferate.

**Fig. 1.**
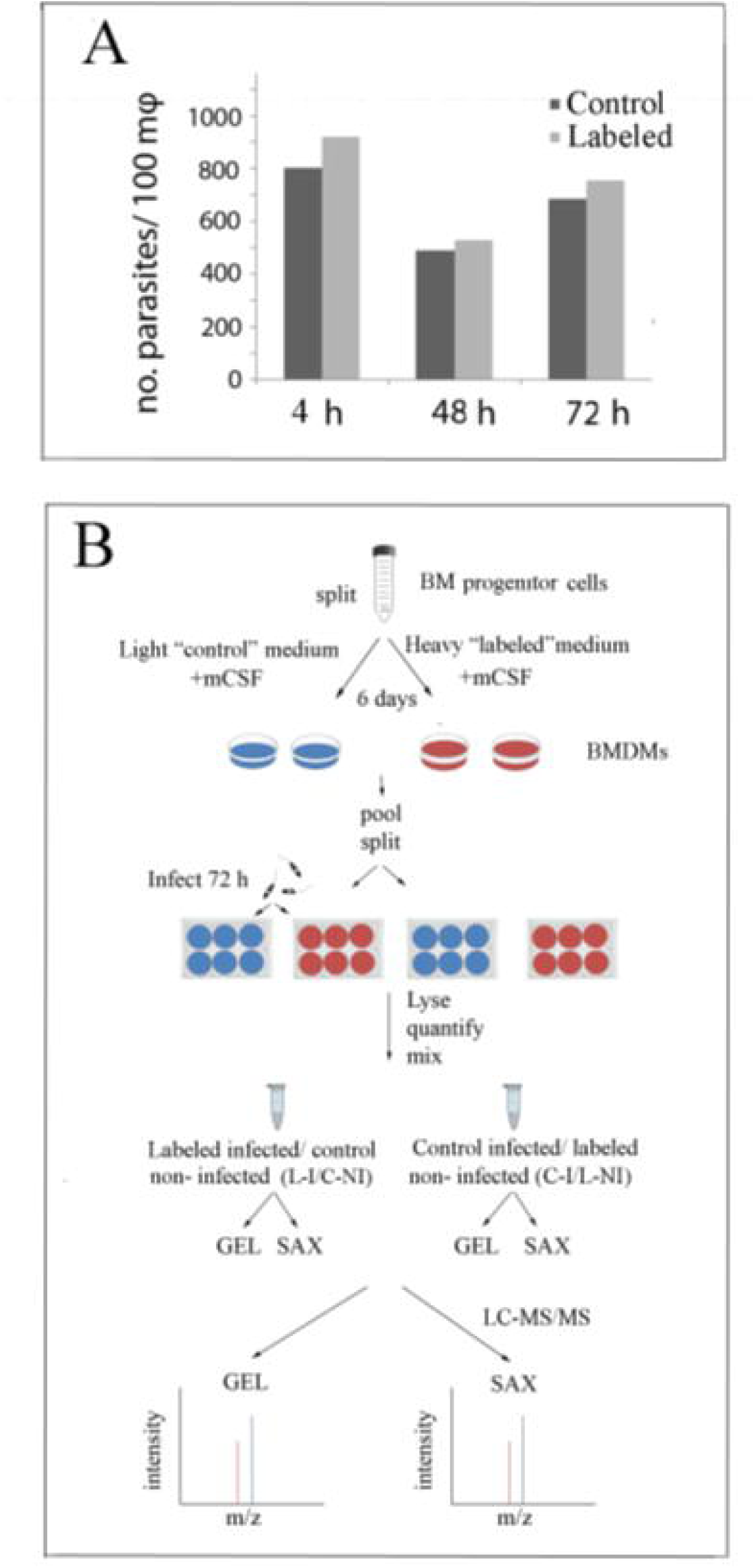
*In vitro* macrophage infection model with *L. donovani* promastigotes and SILAC-based macrophage proteomics analysis. (A) Histogram plots showing the number of intracellular parasites per labeled and control macrophages after 4 h, 48 h and 72 h of infection. (B) Workflow diagram showing the experimental strategy used to reveal the variable proteome of *L. donovani* infected BMDMs. Initially, an equal number of bone marrow (BM) progenitors were cultured and differentiated in the presence of natural amino acids (light “control” medium) or amino acid with heavy isotopes (heavy “labeled” medium) supplemented with 75 ng/mL mCSF. After 6 days of differentiation, adherent cells were detached and plated with fresh control or labeled medium supplemented with 25 ng/mL mCSF. Labeled (red) and control (blue) macrophages were infected or not with stationary phase *L. donovani* promastigotes at a ratio of 10:1 for 4 h. Macrophages were lysed 72 h post-infection, protein extracts were quantified and mixed at a 1:1 ratio in pairs (control vs labeled, infected or not), fractionated by either polyacrylamide gel electrophoresis fractionation (GEL) or strong anion exchange fractionation (SAX), and processed by LC-MS/MS analysis.

In a series of control experiments, we assessed the culture system for variations in host cell phenotype, SILAC label incorporation, infection efficiency and protein fractionation technique. Under our experimental conditions (see Materials and Methods), over 90% of the adherent cells differentiated into a homogeneous macrophage population as judged (i) by expression of the macrophage-specific F4/80 surface antigen as determined by fluorescent microscopy [45, 46], and (ii) the ability of the cells to phagocytose fluorescently labeled, yeast zymosan bioparticles (data not shown). We tested the isotope incorporation efficiency for the amino acids lysine and arginine using LC-MS/MS and established day 6 after the differentiation process, as the minimal culture period to achieve the required incorporation levels of above 95% (data not shown). As shown by the experimental layout in **Fig. 1B**, we performed label-swap experiments, where the experimental state and stable isotope labels are interchanged. Label swap enriches the list of proteins whose abundance is related to the experimental state as it allows for the identification of systematic errors due to labeling, and to experimental outliers. Bone marrow progenitor cells were differentiated into macrophages in medium supplemented with the ‘heavy’ amino acid isotopes, referred to as ‘labelled” (L), and medium supplemented with the ‘light’, natural amino acids, referred to as ‘control’ (C). Cells were infected six days post-differentiation with *L. donovani* promastigotes for 72 h. *L. donovani*-infected (I) and non-infected (NI), labeled (L) and control (C) macrophages were lysed, mixed in equal amounts and their proteomes were analysed by LC/MS-MS. Our protocol further included two fractionation methods, SAX and GEL fractionation that are based on charge- or size-dependent separation, respectively. The reduction of extract complexity by these methodsincreasing the number of quantifications and thus the quality of our proteomics analyses by increasing protein sequence coverage, which facilitates detection of low-abundance proteins.

### Analysis and validation of proteomics data

The comparative analysis of macrophage proteomes from reciprocal L-I/ C-NI and C-I/L-NI samples allowed the identification of 5322 and 5101 proteins using SAX and GEL fractionation, respectively and a combined total 6189 proteins (**Fig. 2A** and **Fig. 2B**). Seventy five percent of proteins identified by GEL and SAX fractionation (3871and 4026 proteins, respectively) were shared between L-I/C-NI and C-I /L-NI (**Fig. 2A**), indicating a very good, qualitative reproducibility in the depth of our analysis. Moreover, the overlap between SAX and GEL was also significant, with sixty five percent 3291 proteins of the proteins identified were shared between the two groups (GEL, SAX: L-I/C-NI and GEL, SAX: L-NI/C-I) (**Fig. 2B**). Volcano plots using GEL and SAX fractionation revealed that a subset of proteins were differentially expressed between in L-I/C-NI and C-I/L-NI (**Fig.1B**).

**Fig. 2.**
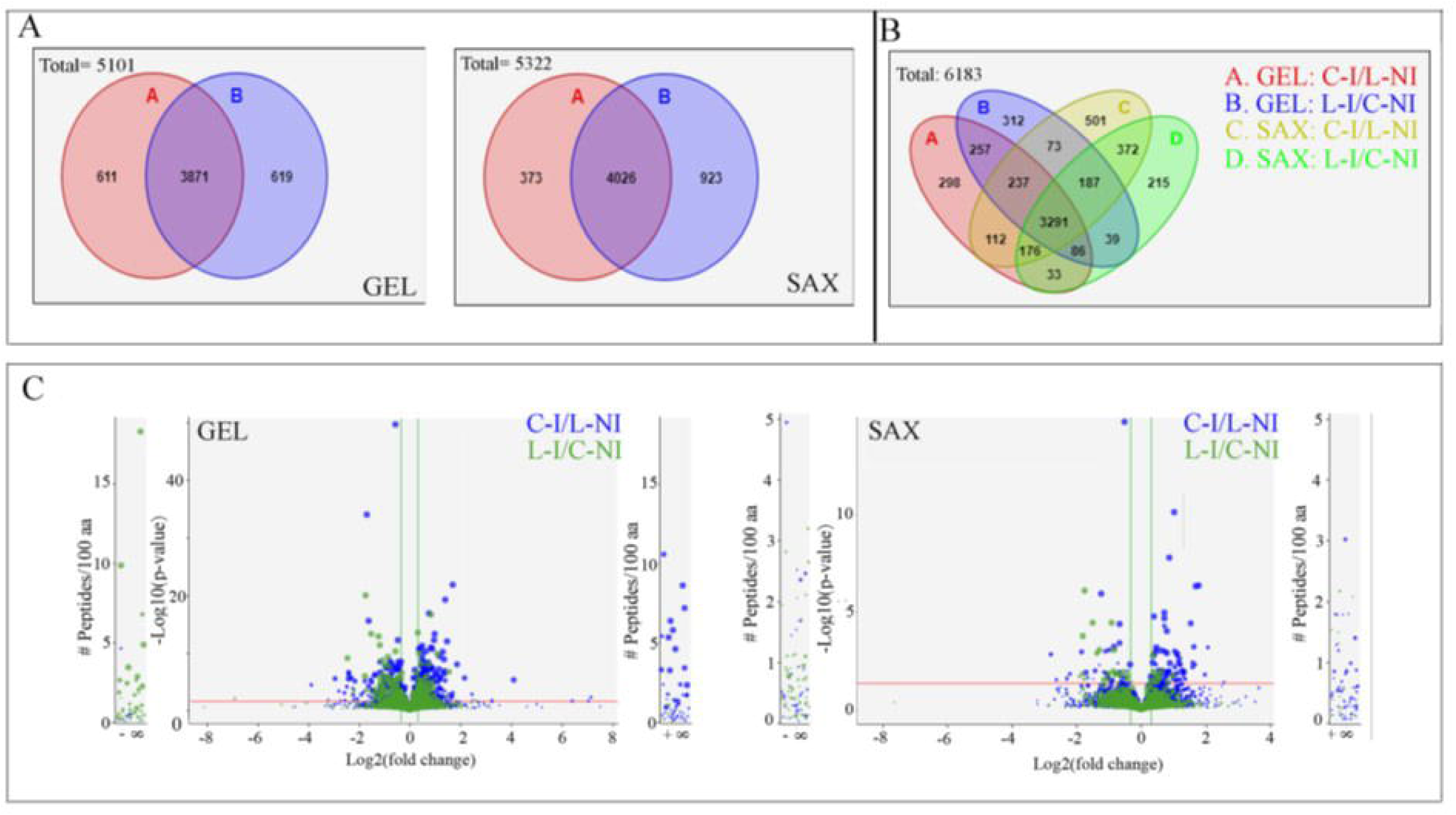
Summary of proteomics results. (A) Venn diagram showing the overlap of identified proteins between L-I/C-NI and C-I/L-NI after GEL (left panel) or SAX fractionation (right panel). (B) Venn diagram showing the combined overlap identified proteins of L-I/C-NI, C-I/L-NI samples in both GEL and SAX fractionation conditions. (C) Volcano plots of the data presented in (A) showing the fold change (FC, x-axis, log2) of abundance for proteins with at least two identified peptides plotted against the p-value (y-axis, −log10). Red vertical and green horizontal lines reflect the filtering criteria, i.e. FC≥1.25 or FC≤0.8, and significance level of 0.05, respectively.

Amongst the proteins with modulated abundance, only macrophage proteins identified in reciprocal (label swapped) experiments that conformed to filtering criteria raised, were selected. To this end, we considered in both forward and reverse experiments a fold change (FC) of 1.25 or higher, with at least 2 peptides identified per quantification. With the exception of infinite quantifications, only quantifications with confidence level of 0.05 or below (p≤0.05), in at least one of the experiments were accepted (**Table 1** and **Table 2**).

This analysis revealed increased abundance in infected macrophages for 35 proteins (27 proteins with GEL, 8 proteins with SAX), with Glutathione S Transferase mu1 (Gstm1/P10649) common in the datasets (**Table 1**). The same analysis revealed reduced abundance for 51 proteins in infected macrophages (42 proteins with GEL, 13 proteins with SAX), with 4 proteins common in the two datasets [the macrophage-expressed gene 1 protein (Mpeg-1/A1L314), Palmitoyl-protein thioesterase 1 (Ppt1/O88531), Cathespin D (Ctsd/ P18242) and the Mannose receptor C-type 1 (Mrc-1/Q61830)] (**Table 2**). Thus, for the primary macrophage proteome, these results indicate that SAX and GEL fractionation procedures together increase the level of quantifications that pass the filtering criteria, increasing the depth of protein quantification.

We then validated our results by immunoblot analysis of infected and non-infected macrophages using β-αctin for normalisation. We confirmed reduced abundance in response to infection for Acid ceramidase 1 (Asah1/Q9WV54), Guanine nucleotide exchange factor for Rab-3A (Rab3IL1, Q8TBN0), Heme oxygenase 1 (Hmox-1/P14901) and Cathepsin D (Ctsd/P18242). Additionally, we confirmed the higher abundance of Glutathione S-transferase Mu 1 (Gstm1/P10649) and Plexin A1 (PlxnA1/P70206) (**Fig. 3A**, **Table 1** and **Table 2**). Total RNA of replica samples used for immunoblot analysis was extracted, and reverse transcription (RT) qPCR was performed for selected transcripts. The transcripts of 60S ribosomal protein L19 (RpL19/P84099) and 14-3-3 protein zeta/delta (Ywhaz/P63101) were used as normalisation controls, as these reference genes are considered to have stable expression [47, 48] in these culture conditions. Certain changes in transcript levels correlated with changes in protein abundance, including reduced abundance for Pppt1 and increased abundance for PlxnA1 and the copper-transporting ATPase 1 (Atp7/Q64430). Other transcripts did not show any correlation, e.g., Ctsd, Rab3IL1 and Hmox-1 mRNAs (**Fig. 3B**). These data suggest that *L. donovani* both affects transcriptional and post-transcriptional regulation of infected BMDMs.

**Fig. 3.**
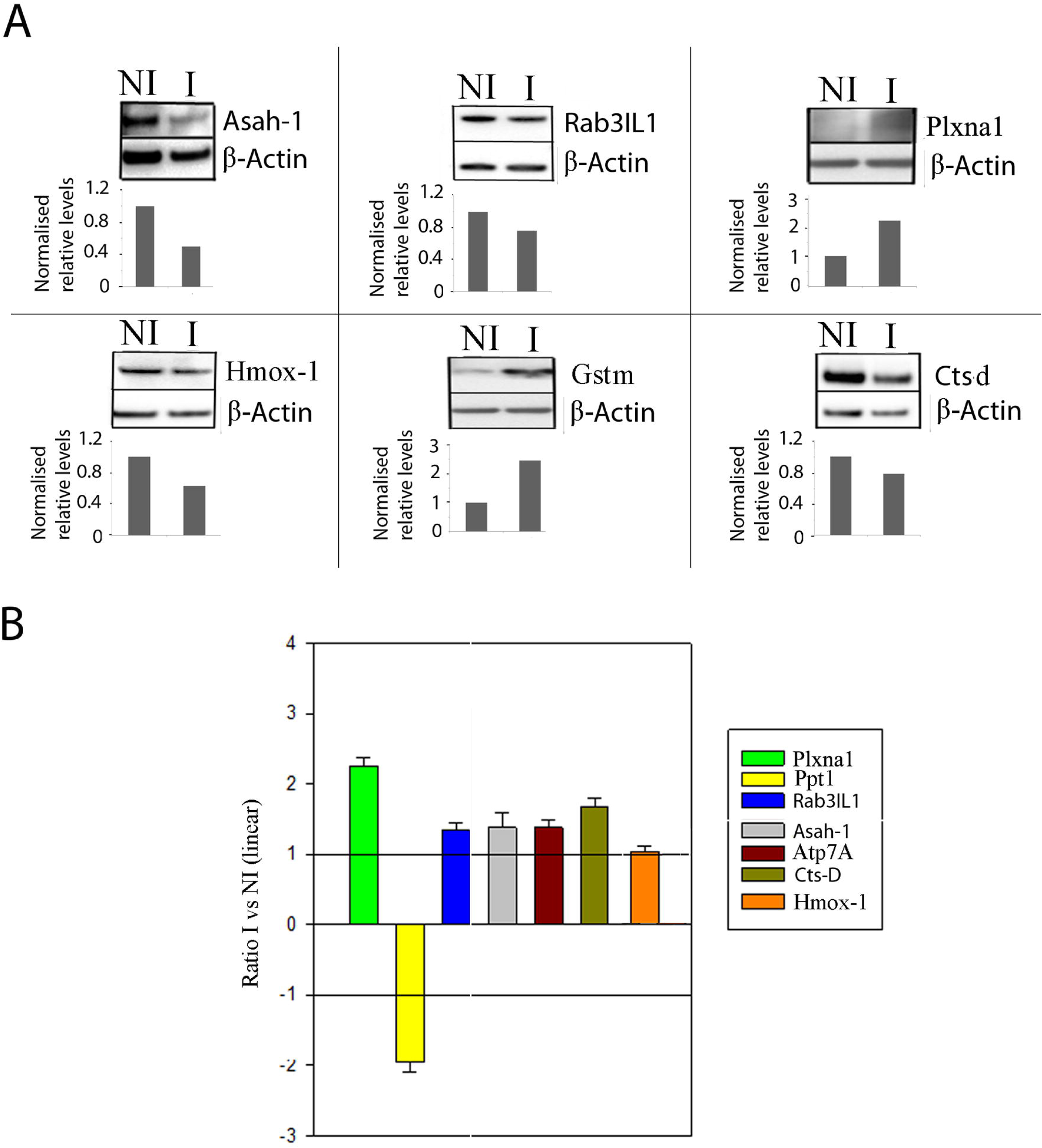
Differential abundance for BMDM proteins and transcripts at 72 h post *L. donovani* infection. (A) Immunoblot analysis. 20 μg of total protein extracts from infected (I) and non-infected (NI) BMDMs were analysed for Asah-1, Rab3IL1, Plxna1, Hmox-1, Gstm, and Ctsd proteins. β-actin expression was used as a loading control. The intensities of the bands were analysed using the Image J software. The fold overexpression, represented as a bar diagram, was calculated by dividing the band intensity representing the protein of interest with the band intensity of β-actin. Results are representative for two independent experiments. (B) RT-qPCR. Column diagrams showing the mean ratio from three different experiments of RNA abundance in infected versus non-infected macrophages for Plxna1, Ppt1, Rab3IL1, Asah-1, Atp-7A and Hmox-1 relative to Ywhaz and RpL19. Error bars represent the standard deviations of three experiments.

### Gene Ontology analysis illuminates the landscape of subverted pathways in *L. donovani* infected macrophages

The datasets of differentially modulated proteins were analysed for enriched biological processes, molecular functions and cellular components using the Gene Ontology (GO) and the *Mus musculus* Uniprot databases (https://www.uniprot.org/proteomes/UP000000589), and results were visualised with the BinGO plug-in of the Cytoscape 3.5.1 software package [43]. Up-regulated biological processes were related to metabolism, including lipopolysaccharide biosynthetic process (GO-ID: 9103), cellular ketone metabolic process (GO-ID: 42180), oxoacid (GO-ID: 43436) and carboxylic acid (GO-ID: 19752) metabolism (**Fig. 4A**, **Table S2**), the latter enrichment having previously been observed in THP-1 cells infected with *L. donovani* promastigotes [28]. Enrichment was also observed for various molecular functions, including intramolecular oxidoreductase activity transposing S-S bonds (GO-ID: 16864), intramolecular oxidoreductase activity, interconverting keto and enol groups (GO-ID: 16862) and protein disulfide isomerase activity (GO-ID: 3756)(**Fig. 4B, Table S2**). This last activity is important for protein folding, stress response (balancing the effects of oxidative stress) and phagocytosis [49]. Moreover, enrichment in cellular components included the cytoplasm (GO-ID: 5737), intracellular membrane bounded organelle (GO-ID: 43231) and endoplasmic reticulum (GO-ID: 5783), with many of the up-regulated endoplasmic reticulum (ER) proteins involved in protein folding and post-translation modification (**Fig. 4C**, **Table S2**), revealing a potential impact of *L. donovani* infection on endoplasmic reticulum stress and the unfolded protein response as previously documented [50].

**Fig. 4.**
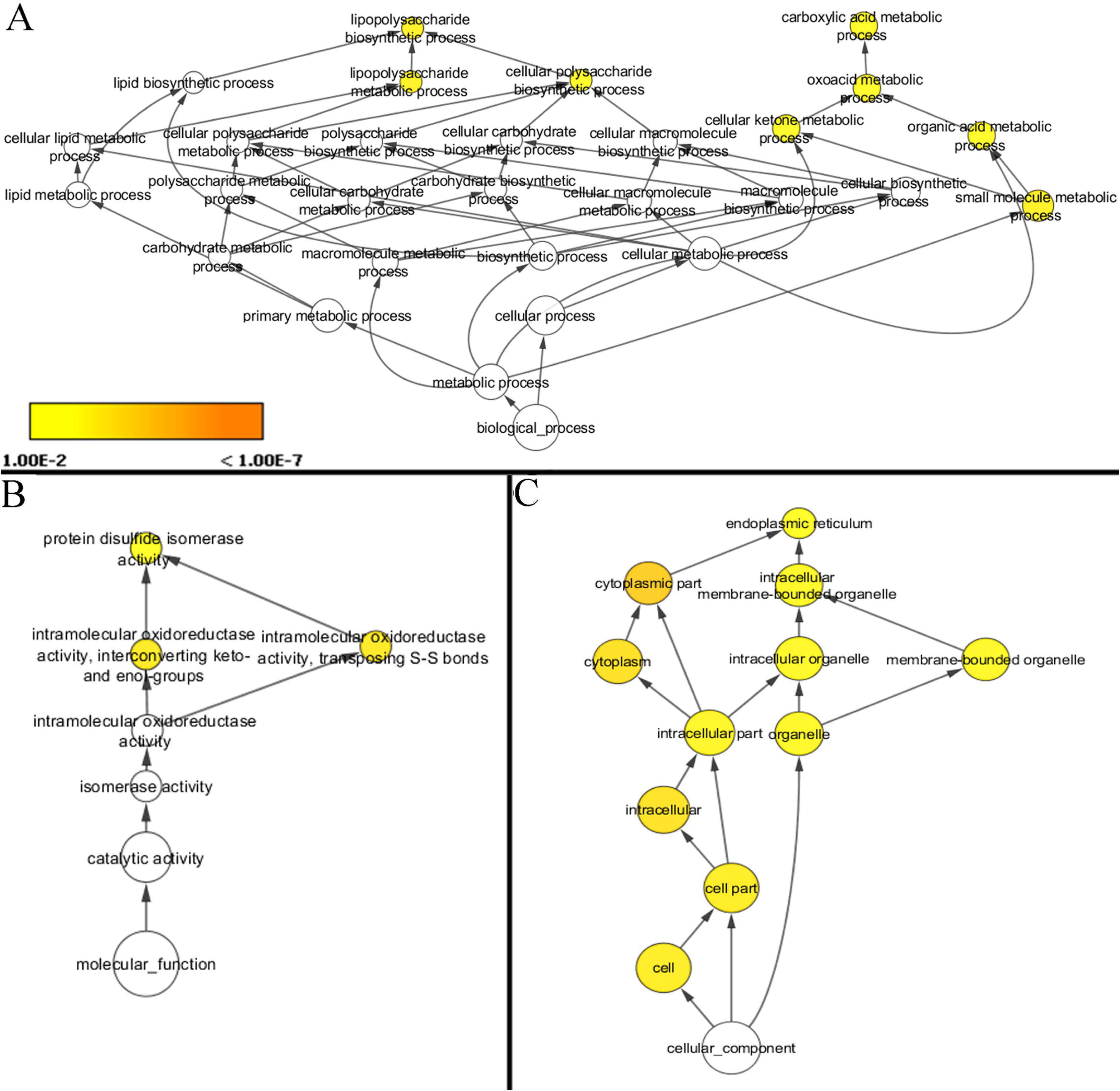
Gene Ontology analyses for up-regulated proteins in *L. donovani* infected BMDMs. (A) GO analysis for Biological processes. (B) GO analysis for Molecular functions. (C) GO analysis for Cellular component. The analyses were performed using the hypergeometric statistical test, a Benjamini & Hochberg false discovery rate and significance level of 0.01. The p-value is indicated by the color according to the legend.

Using the data set of proteins showing reduced abundance, enrichment was observed for biological processes involved in vacuole (GO-ID: 7033) and lysosome (GO-ID: 7040) organisation and catabolic process (GO-ID: 9056) (**Fig. 5A, Table S3**). Enriched molecular functions in the down-modulated dataset, included binding (GO-ID-5488), transcription elongation regulator activity (GO-ID: 3711), hydrolase activity hydrolysing o-glycosyl compounds (GO-ID: 4553) and cysteine type endopeptidase activity enriched in cysteine type endopeptidases (GO-ID: 4197) (**Fig. 5B, Table S3**). Moreover, amongst the enriched cellular components were lytic vacuole (GO-ID: 323), lysosome (GO-ID: 5764) and vacuole (GO-ID: 5773), and thus the very organelles and structures exploited by the parasite as intracellular niche. Other enriched cellular components included cytoplasm (GO-ID: 5737), intracellular membrane bounded organelle, (GO-ID: 43231) and integrin complex (GO-ID: 43231) (**Fig. 5C, Table S3)**.

**Fig. 5.**
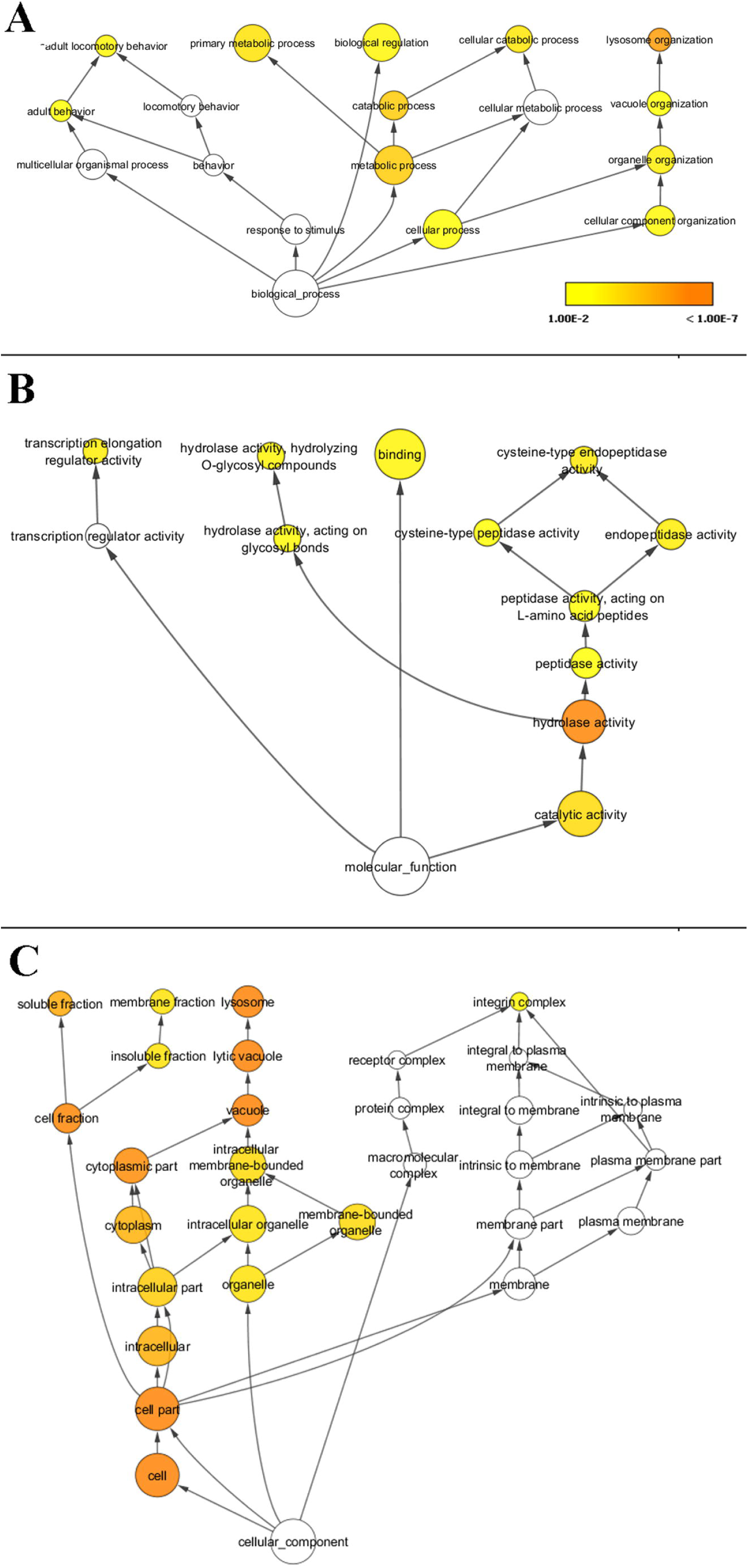
Gene Ontology analyses for down-regulated proteins in *L. donovani* infected BMDMs. (A) GO analysis for Biological processes. (B) GO analysis for Molecular functions. (C) GO analysis for Cellular component. The analyses were performed using the hypergeometric statistical test, a Benjamini & Hochberg false discovery rate and significance level of 0.01. The p-value is indicated by the color according to the legend.

### *L. donovani* modulate the abundance of proteins involved in the lysosome and lytic vacuole organisation of BMDMs

Our study reveals that *L. donovani* promastigote infection has a profound effect on BMDM lysosomal biology, with remodeling of this organelle likely promoting parasite survival in its known intracellular niche, the phago-lysosome [4, 51]. Our data show that the ratios of a total of 15 lysosomal proteins in both forward [L-I/C-NI, ratio 1, (r1)] and reverse experiments [C-I/L-NI, ratio 2, (r2)] were below 0.8 (**Table 2** and **Table S3**). Down-regulated proteins included various hydrolases such as lysosomal cathepsins (Ctsb/P10605, r_1_ =0.59 r_2_=0.54; Ctsc/P97821, r_1_=0.49, r_2_=0.41; Ctsd/P18242, r_1_=0.35, r_2_=0.59) or dipeptyl peptidase 2 (Dpp-2/Q9ET22, r_1_=0.4, r_2_=0.75) and the Lysosome-associated membrane glycoproteins, Lamp2 (Cd107b/P17047, r_1_=0.72, r_2_=0.7) and Lamp4 (Cd68/P31996, r_1_=0.68, r_2_=0.49) (**Table 2**).

Lysosomal proteases, including cathepsins may affect parasite survival as they carry microbicidal activity and are key regulators of the host immune response by modulating Major histocompatibility Class (MHC) II antigen presentation and the trafficking of <u>T</u>oll like receptors (TLRs) [52]. Down-modulation of different cathepsins may cause distinct effects on macrophage activation. For example, deletion or pharmacological inhibition of Ctsb results in inflammatory phenotypes [53, 54] and Ctsd inhibition blocks both Th1 and Th2 responses by strongly suppressing MHCII-associated invariant chain molecule [54]. Different *Leishmania* species and strains, parasite stages (promastigotes/amastigotes) and host species may modulate cathepsins in different ways. For example, Prina *et al* reported a time-dependent increase of Ctsb, Ctsh and Ctsd activity in rat bone marrow-derived macrophages infected with *L. amazonensis* [55]. On the other hand, the RNA transcripts of Ctsc and Ctss were reported to be down-modulated in monocyte derived human macrophages 24 h after *L. major* metacyclic promastigote infection [56].

Apart from lysosomal peptidases, a number of hydrolases involved in fatty acid metabolism were also affected in macrophages by *L. donovani* infection. Interestingly, the down-modulation of the lyase Palmitoyl-protein thioesterase 1 (Ppt1/O88531, r_1_=0.23, r_2_=0.3) involved in the removal of thioester-linked fatty acyl groups (like palmitate) is anticipated to disrupt lysosomal catabolism via the rapid accumulation of palmitoylated proteins [57]. Moreover, Ppt1 down-modulation impairs mTOR signaling [57] the repression of which has been also linked to *Leishmania* intracellular survival [58]. Additionally, down-modulation of Acid ceramidase −1 (Asah-1/Q9WV54, r_1_**=** 0.45, r_2_=0.5) (**Table 2**) - an enzyme that hydrolyses the sphingolipid ceramide into sphingosine and free fatty acid - may also impact the macrophage environment to favor parasite survival. *Leishmania* is favored by increase in ceramide concentration and in turn *Leishmania* infection, modulates the up-regulation of ceramide synthesis in a biphasic way [59]. At early time points after *Leishmania* infection, increase is mediated by Acid Sphingomyelinase activation, which catalyses the formation of ceramide from sphingomyelin, promoting the formation of cholesterol rich lipid microdomains [59], required for *L. donovani* uptake by macrophages [60]. In the second phase, the *de novo* biosynthesis of ceramide is induced via ceramide synthase, resulting in the displacement of cholesterol from the membrane and impacting MHC-II class antigen presentation [59]. Moreover, ceramide increase leads to severe host immune suppression via the reduction of nuclear translocation of NFkB and AP-1, the up-regulation of immunosuppressive cytokines TGF-β and IL-10, and reduced NO generation [61]. Thus increase in ceramide concentration is required for parasite survival, and the down-modulation of Asah-1 revealed in this screen, could be an additional mechanism contributing to this rise, and hence merits further investigation.

Overall these results suggest a major modification of lysosomal components that might aid the parasite to survive inside the hostile macrophage environment. The mechanism that contributes to lysosomal protein down-modulation after *L. donovani* infection, is elusive. One possible scenario would be a dysregulation of transcription factor EB, which is a master regulator of lysosomal biogenesis that mediates the transcription of many lysosomal hydrolases [62]. Another scenario would be that the lysosomal content is modified via the action of lysosomal exocytosis, which takes place to repair host cell plasma wounding caused by the parasite [63] or by the direct interaction of the parasites’ exo-proteases with lysosomal proteins, leading to protein degradation.

### *L. donovani* induce alterations in abundance of macrophage proteins involved in immunomodulation

Our data demonstrate that *L. donovani* promastigotes inhibit host cell immune functions by down-regulation of several macrophage proteins that participate in immunomodulation and innate immunity (Cd180/Q62192, r_1_=0.66, r_2_=0.76; Tlr13/Q6R5N8, r_1_=0.66, r_2_=0.5; Grn/P28798, r_1_=0.7, r_2_=0.62; Arrb2/Q91YI4, r_1_=0.61, r_2_=0.4; Gpr84/Q8CIM5, r_1_=0.64 and r_2_=1/∞; Hmox-1/P14901, r_1_=0.67, r_2_=0.39; Mpeg1/A1L314, r_1_=0.31, r_2_=0.4) (**Table 2**). These results indicate that infection likely dampens the macrophage pro-inflammatory response given the observed down-modulation of (i) Tlr13, which acts via Myd88 and Traf6 leading to NF-κΒ activation [64], (ii) Granulin (Grn) a soluble cofactor for Tlr9 signaling [65], and (iii) the G-protein coupled receptor Grp84 known to be activated by medium-chain free fatty acids to enhance inflammation and phagocytosis in macrophages [66].

Moreover, *L. donovani* promastigote infection results in increased abundance of macrophage proteins that act as markers or as regulators of alternatively activated macrophages (M2-like phenotype) (**Table 1**) in agreement with previous reports [67–72]. Amongst these proteins were: (i) the Cd9 antigen (P40240, r_1_=1.41, r_2_=1.76), a tetraspanin family member known to form a complex with integrins and to negatively regulate LPS-induced macrophage activation [73], (ii) the highly up-regulated urokinase-type plasminogen activator (Plau/P06869, r_1_=6.02, r_2_=5.41) known to polarise macrophages towards a M2-like phenotype [74], (iii) the monoamine oxidase A (Mao-A/Q64133, r_1_=1.31, r_2_=2.24) involved in the breakdown of monoamines [75], (iv) the TNF alpha induced protein 8 like 2 (Tnfaip8l2/Q9D8Y7, r_1=_ ∞ and r_2_=1.4) involved in phospholipid metabolism [76, 77], (v) the Cd93 antigen (Cd93/O89103, r_1_=1.35, r_2_=1.76), a C-type lectin transmembrane receptor [78] and (vi) the fatty acid-binding protein (Fabp4/P04117, r_1_=1.3, r_2_=1.57) that delivers long-chain fatty acids and retinoic acid to their cognate receptors in the nucleus [79] (**Table 1**). Interestingly, Fabp4 up-modulation in infected macrophages is in line with other studies after macrophage infection with *L. major* promastigotes [26] and *L. amazonensis* amastigotes [24], and suggest that the increased abundance of this protein maybe independent of the *Leishmania* stage or species.

In contrast to proteins that control, or are enriched in alternatively activated macrophages, two proteins less abundant in infected macrophages, Cd180 and Beta-arrestin-2 (Arrb2), are known to either promote or reduce inflammation depending on the cell type and infection system [80–83]. Moreover a protein known to be enhanced in alternatively activated macrophages, Hmox-1 that converts pro-oxidant heme to the antioxidant biliverdin and bilirubin, restoring the redox environment [84] was found down-modulated in this screen in response to *Leishmania* infection (**Table 2**). Other studies have shown that Hmox-1 is up-regulated in mouse peritoneal macrophages infected with *L. infantum (L. chagasi*) promastigotes and promotes the persistence of the parasite [85]. In our infection system, despite the repressed protein levels, we find Hmox-1 mRNA to be up-regulated in macrophages infected with *L. donovani*. The negative correlation between Hmox-1 RNA and protein levels, amongst other factors, could be due to increased secretion of Hmox-1 in the culture medium, as Hmox-1 can be a secreted protein [86].

### *L. donovani* up-modulate the abundance of proteins involved in the defense against oxidative and ER stress in BMDMs

Proteins related to oxidative stress and ER stress response were more abundant in BMDMs 72 h after *L. donovani* infection, including proteins involved in glutathione metabolism, such as Glutamate-cysteine ligase regulatory subunit (Gclm/O09172, r_1_=1.28, r_2_=1.29), the Glutathione transferase mu1 (Gstm1/P10649, r_1_=1.36; r_2_=2.53), ferritin light (Ftl1/ P29391, r_1_=2.9, r_2_=2.64) and heavy (Fth1/P09528, r_1_= 1.37, r_2_=2.23) chains, and BolA like 1 (Bola1/Q9D8S9, r_1_=1.73 and r_2_=∞), a protein that prevents mitochondrial changes upon glutathione depletion [87] acting as mitochondrial iron-sulfur cluster assembly factor [88] (**Table 1**). All of these proteins are known to normalise the redox state of the cell or protect the cell from oxidants [87, 89–91]. Interestingly, up-regulated transcripts involved in the response against oxidative stress and glutathione metabolism were previously shown to be up-regulated in human monocyte derived macrophages infected with *L. (Vianna) panamensis* promastigotes [9] and in mouse BMDMs infected with *L. major* promastigotes [26]. Moreover, up rise in cellular ROS, could be associated with ER stress, the unfolded protein (UPR) response and a concomitant up rise of ER proteins aiding protein folding, as a mechanism of the cell to repair oxidant damage [92]. In our system several of these proteins including disulfide-isomerase forms [Protein disulfide-isomerase A3 (Pdia3/ P27773, r_1_=1.39,r_2_=1.27), A4 (Pdia4/P08003, r_1_=1.35, r_2_=1.27) and A6 (Pdia6/Q922R8, r_1_=1.43 r_2_=1.31)] [93] and Hypoxia up-regulated 1 (Hyou1/Q9JKR6, r_1_= 1.86, r_2_=1.38 [94], displayed higher levels in infected macrophages (**Table 1**). Likewise, ER stress and induction of the unfolded protein response was previously documented for RAW 264.7 cells infected with *L. amazonensis* [50]. Overall, these results suggest that *L. donovani* infection induces ROS production and ER stress [10, 95], with specific proteins involved in this process displaying higher abundance, and likely acting as a defense mechanism against the oxidants’ deleterious effects.

### *L. donovani* modulates the abundance macrophage proteins involved in intracellular trafficking and ion movement

The abundance of many macrophage proteins involved in intracellular trafficking was modified in *L. donovani* infected BMDMs. Previous studies have shown that *Leishmania* infection impairs intracellular trafficking by modulating small GTPase signal transduction [28]. This form of signaling is important in host/pathogen interactions as it regulates pathways involved in phagocytosis and oxidative burst, vesicle fusion and actin organisation [96, 97]. The deregulation of small GTPase signal transduction, likely alters these macrophage functions, and produces a favorable environment for intracellular parasite infection [28, 98, 99]. Our screening extends previous studies and provides evidence of altered protein expression data in proteins involved in small GTPase signalling, with four proteins showing reduced abundance during infection (Gmip Q6PGG2, r_1_=0.79, r_2_=0.49; Rgl2/Q61193, r_1_=1/∞; r_2_=0.35; Arfip2/Q8K221, r_1_=1/∞ r_2_=0.64; Rab3il1/Q8VDV3, r_1_=0.66, r_2_=0.39; Nisch/Q80TM9, r_1_= 0.66, r_2_=0.46) (**Table 2**), and one member showing increased abundance (Tbc1d22a/Q8R5A6, r_1_=∞ and r_2_=1.89) (**Table 1**). Moreover, various other proteins affecting intracellular trafficking were modulated in response to *L. donovani* promastigote infection, including a signal recognition particle receptor subunit beta (Srprb/P47758; r_1_=1.36, r_2_=1.25) (**Table 1**) that likely restores trafficking imbalances to the ER [100], and two cytoskeletal organisation proteins (Peak1/Q69Z38, r_1_=0.45, r_2_=0.46; Ppp2r5e/Q61151, r_1_=1/∞, r_2_=0.73) [101, 102]. Additionally, the abundance of macrophage mannose receptor 1 (Mrc-1) – a protein involved in endocytosis and phagocytosis - was down-modulated (Q61830, r_1_=0.63, r_2_=0.33), in agreement with previous studies performed with mouse peritoneal macrophages infected with *L. donovani* promastigotes [103]. With the exception of Mrc-1, the majority of these macrophage proteins have not been previously shown to be modulated by *Leishmania* infection. The modulation of these proteins merits further investigation, as they could be part of the parasite evasion strategy interfering with vesicular fusion events in the host cell, important for parasite survival.

Interestingly, another form of trafficking, ion movement, has been associated with inflammatory response [104–106]. For example, zinc/copper imbalance reflects immune dysregulation in human leishmaniasis [106]. In this study, some proteins involved in ion transport were modulated in infected BMDMs, including (i) Atp1a1 (Q8VDN2, r_1_= 1.39; r_2_=1.26) a subunit of the sodium potassium ATPase [107](**Table 1**), (ii) stomatin (Stom/P54116, r_1_= 0.78, r_2_=0.48) (**Table 2**) known to regulate ion channel activity and transmembrane ion transport [108] and (iii) copper-transporting ATPase 1 (ATP7a/P56542, r_1_=2.61, r_2_=2.66) that regulates copper efflux under conditions of elevated extracellular copper concentration [109] (**Table 1**). Overall, these changes may reflect immune function and their influence on host/pathogen interaction merits further investigation.

### *L. donovani* modulate the abundance of extracellular matrix and adhesion proteins in BMDMs

Macrophage adhesion and cell-cell interactions with other immune cells play important roles in health and disease, with co-stimulatory molecules regulating T cell activation or cell migration enabling leukocyte recruitment to sites of inflammation. Our proteomics screen revealed changes in expression of extracellular matrix and adhesion proteins that could explain the reported modified adhesion of infected macrophages to connective tissue [110]. We confirmed down-modulation of the co-stimulatory molecule Itgax (Q9QXH4, r_1_=1/ ∞, −r_2_= 0.33) (**Table 2**). Changes in abundance of the integrin subunit Itgβ5 (O70309, r_1_= 0.73 and r_2_=0.55) which acts with Itga5 as a fibronectin receptor, were detected, as well as differences in other adhesion or adhesion signaling molecules (Nisch/Q80TM9, r_1_=0.66, r_2_=0.56; Plxna1/ P70206, r_1_= 1.65, r_2_=2.46; Lpxn/Q99N69, r_1_= 0.74, r_2_=0.65) (**Table 1** and **Table 2**). Thus, our data expand and further emphasise the impact of intracellular *Leishmania* infection on connective tissue remodeling that may modify host adaptive immune responses mediated by infected macrophages [110, 111].

### *L. donovani* modulate the abundance of macrophage proteins involved in gene expression

It is well established that *Leishmania* infection modulates macrophage gene expression [21]. In our study, several proteins implicated at different levels of gene expression regulation, showed differential abundance in response to *L. donovani* BMDM infection, including transcriptional, epigenetic or post-transcriptional regulators (Hist4h4/P62806, r_1_=1.44, r_2_=1.33; Lpxn/Q99N69, r_1_= 0.75, r_2_=0.65; Ddx42/ Q810A7, r_1_=0.74, r_2_=0.65; Psmc3/088685, r_1_=1.34, r_2_=1.25; Supt5h,/O5520, r_1_= 0.67; r_2_=0.68; Supt6h/Q62383, r_1_=0.59, r_2_=0.64), proteins that modify RNA (Cmtr1/Q9DBC3, r_1_=1.39 and r_2_=∞; Dus3L/Q91XI1, r_1_= 0.34, r_2_=1/∞), as well as translation regulators (eIF-3/Q9QZD9, r_1_= 0.77, r_2_=0.73; Rps6ka1-Rsk1/P18653, r_1=_0.43, r_2_=1/∞) (**Table 1** and **Table 2**). The higher abundance of histone H4 (Hist4h4) in infected macrophages is in accordance with previous reports for *L. donovani* infected THP-1 cells [28], whereas up-regulation of histones is anticipated to have a negative effect on gene expression by the modulation of chromatin structure [112]. Moreover, the lower levels of focal adhesion regulator Lpxn [113, 114] and the increased abundance of the 26S proteasome ATPase component, Psmsc3/TBP-1 [115], two proteins that display novel functions and are known to modulate transcription via specific nuclear receptors [116–118]. Thus the modulation of abundance of these proteins in infected macrophages, may affect transcriptional regulation of *Leishmania* infected macrophages. In addition, Psmsc3/TBP-1 regulates the transcription of Class II trans-activator (CIITA), the master regulator of the MHC-II transcription complex and a critical factor for the initiation of the adaptive immune response [119]. In addition, the down-modulation of specific transcriptional elongation factors (Supt5h and Supt6h) [120–122] is anticipated to contribute to transcriptomic changes in infected macrophages. The down-modulation of the eukaryotic translation initiation factor eIF-3 and of protein S6 kinase alpha 1(Rps6ka1/Rsk1) - a kinase known to stimulate protein synthesis and a key mediator of mTOR function [123] - suggests that protein translation proceeds at lower rates in infected macrophages [123]. Rps6ka1/Rsk1 is known to “connect the stress-induced activation of transcription factors and mitogens to the ribosome” [124]. This connection is mediated by the activation/phosphorylation of c-Fos, IκBα, cAMP-response element-binding protein (CREB) and CREB-binding protein [125], proteins involved in the inflammatory response. Thus, the modulation of this kinase likely participates in the reprogramming of *Leishmania-*infected macrophages to favor intracellular parasite survival.

### L. donovani modulate the abundance of macrophage proteins involved in metabolic processes

Previous studies have shown that *Leishmania* parasites exploit host metabolism to survive inside the hostile macrophage environment. It has been demonstrated that, early during *Leishmania* infection, the aerobic glycolysis predominates with inhibition of the TCA cycle [26], whereas at later time points energy metabolism is shifted towards oxidative phosphorylation and TCA cycle [28, 126]. *Leishmania* infection also perturbs lipid metabolism, including sterol biosynthesis and triacylglyceride synthesis in BMDMs [24, 26]. In our screen, the cellular ketone metabolic process (GO-ID: 42180) and oxoacid metabolic process (GO-ID: 43436) - a broad metabolic processes that includes TCA cycle and aminoacid biosynthesis, were modified, with 5 proteins showing increased abundance (Fabp4/P04117, r_1_= 1.3, r_2_=1.57; Sucla2/Q9Z2I9, r_1_=1.27, r_2_=1.37; Phyh/O35386, r_1_=∞ and r_2_=1.25; Cmas/Q99KK2; r_1_= 1.27, r_2_=1.72; Gclm/O09172, r_1_=1.27, r_2_=1.28). Indeed, Succinate--CoA ligase [ADP-forming] subunit beta (Sucla2) – an enzyme that couples the hydrolysis of succinyl-CoA to the synthesis of ATP [127] - was one of the TCA cycle proteins shown to be up-regulated in *L. donovani*-infected THP-1 cells [28]. Likewise, Sterol O-acyltransferase 1 (Soat1/Q61263, r_1_=1.29, r_2_=1.31) - a cholesterol metabolism enzyme known to form cholesteryl esters from cholesterol [128] – displayed higher levels in infected macrophages (**Table 1**). This change could be related to the increase in cholesterol biosynthesis of macrophages infected with *Leishmania* [24, 26].

Finally, fatty acid metabolism, including fatty acid elongation (Ppt1/ O88531, r_1_= 0.23, r_2_=0.3), phospholipases of the ether lipid metabolic process [(Pla2g7/Q60963, r_1_= 0.68, r_2_=0.35; Plbd2/ Q3TCN2, r_1_=0.71, r_2_=0.49) and hydrolases of the sphingolipid metabolism (Asah-1/Q9WV54, r_1_=0.45, r_2_=0.5; Gla/P51569, r_1_=0.63, r_2_=0.75; Glb1/P23780, r_1_=0.55, r_2_=0.76), were found to be less abundant in infected macrophages (**Table 2**). Fatty acid metabolism, including sphingolipid metabolism [129] has been shown to be modulated in macrophages infected with *Leishmania* [24, 26, 59] and is likely to play an important function in *Leishmania*/macrophage interaction.

### L. donovani modulate the abundance of macrophage proteins that regulate glycan and glycoside protein post-translational modifications

In addition, proteins regulating protein sugar (glycan and glycoside) post-translational modifications were modulated in *L. donovani* infected macrophages, with decreased levels observed for (i) ribophorin I (RpnI/Q91YQ5, r_1_=1.65, r_2_=1.28), an essential subunit of the N-oligosaccharyl transferase (OST) complex which catalyses the N-glycosylation of proteins [130] (**Table 1**), and (ii) alpha and beta galactosidases (Gla/P51569, r_1_=0.63, r_2_=0.75; Glb1/Q91YQ5, r_1_=0.55, r_2_=0.76). In contrast, increased abundance was observed for cytidine monophospho-N-acetylneuraminic acid synthetase (Cmas/Q99KK2, r_1_=1.27, r_2_=1.72) that catalyses the activation of N-acetylneuraminic acid (NeuNAc) to cytidine 5’-monophosphate N-acetylneuraminic acid (CMP-NeuNAc) [131], a substrate required by sialyltransferases for the addition of sialic acid to growing oligosaccharide chains. Conceivably, changes in protein glycosylation may affect protein function and alter immune responses and cell adhesion signaling [132, 133] in infected macrophages, that may favor parasite survival.

### Conclusions

In conclusion, our analyses establish a novel experimental framework for the quantitative proteomics analysis of *Leishmania* infected primary macrophages. Our results draw a highly complex picture of *Leishmania*/macrophage interaction and highlight the pleiotropic modulation of biological processes and molecular functions in infected BMDMs that likely establish permissive conditions for intracellular parasite survival and chronic infection. The parasite seems to have developed mechanisms to subvert key macrophage functions and trigger changes in host cell metabolism, innate immunity and lysosomal function. Future experimental validation combining systems-level, phenotypic, and functional genetic analyses is required to correlate enrichment/activation or depletion/inhibition of identified host pathways with changes in intracellular *Leishmania* survival.

## Data linking

The mass spectrometry proteomics data have been deposited to the ProteomeXchange Consortium via the PRIDE partner repository [134] with the dataset identifier PXD013448 (username, password: APPQoJX0).

## Acknowledgements

This work was supported by the Institute Pasteur Transverse Research Programs [PTR, grant no.: PTR539], the French Government’s Investissement d’Avenir Program, Labex Integrative Biology of Emerging Infectious Diseases [grant no.: ANR-10-LABX-62-IBEID], and by “Région Ile-de-France” [grant no.: 2013-2-EML-02-ICR-1] and Fondation pour la Recherche Médicale grants [grant no.: DGE20121125630].

