## Supplemental material for "SILAC-based quantitative proteomics reveals pleiotropic, phenotypic modulation in primary murine macrophages infected with the protozoan pathogen *Leishmania donovani*"

**SUPPLEMENTAL FILE**

**Table S1 List of primers used for RT-qPCR analysis**

| **Gene/ Ensemble id** | **Primer sequence 5' 3'** |
| --- | --- |
| **Asah-1** | **F:**ACCTGTCCTCCACAAGC |
| /ENSMUSP00000034000 | **R:** CCAGCCTATACAAGGGTCT |
| **ATP7-A** | **F:**CCCGAGTGATAGCAGAGTTTA |
| /ENSMUST00000113557.7 | **R:**GAAGCACACGTCATTCCT |
| **Ctsd** | **F**:TTCCTTGTGAGAAGGTGTCC |
| /ENSMUSP00000121203 | **R**:CACTCAGGCAGATTGTCTTT |
| **Hmox-1** | **F**:TTTCAGAAGGGTCAGGTGTC |
| /ENSMUST00000005548 | **R**:CTGGGTTCTGCTTGTTGC |
| **Plexin A1** | **F:** TGTCAGTCACCATGTCCC |
| /[ENSMUST00000163139.7](https://www.ensembl.org/Mus_musculus/Transcript/Summary?db=core;g=ENSMUSG00000030084;r=6:89316314-89362620;t=ENSMUST00000163139) | **R:**GTCTCCGTGAAGAGTCCTCAAA |
| **Ppt1** | **F:** TGCCCAGGAGAGAGTTCTCA |
| [ENSMUST00000030412.10](https://www.ensembl.org/Mus_musculus/Transcript/Summary?db=core;g=ENSMUSG00000028657;r=4:122836242-122859175;t=ENSMUST00000030412) | **R:** AGTACTGTGCTTGCACCA |
| **Rab3IL1** | **F:**CCAACATGAAGCAAAGGCGA |
| /[ENSMUST00000121418.7](https://www.ensembl.org/Mus_musculus/Transcript/Summary?db=core;g=ENSMUSG00000024663;r=19:10001669-10038380;t=ENSMUST00000121418) | **R:**GGTGTGGATGTGATGACC |
| **Ywhaz** | **F:**GTTACTTGGCCGAGGT |
| /ENSMUSG00000022285 | **R:**GGAGTTCAGGATCTCGT |
| **RpL19** | **F:**TACTGCCAATGCTCGG |
| /ENSMUSG00000017404 | **R:**AACACATTCCCTTTGACC |

**Table S2 Up-regulated GO categories in infected macrophages with *L. donovani* promastigotes**

|  | BIOLOGICAL PROCESS | | | | | | |
| --- | --- | --- | --- | --- | --- | --- | --- |
| GO-ID | Description | p-val | corr p val | cluster frequency | | total frequency | genes |
| 9103 | lipopolysaccharide biosynthetic process | 1.16E-05 | 2.23E-03 | | 2/26 7.6% | 6/28950 0.0% | Q99KK2 Q8K297 |
| 8653 | lipopolysaccharide metabolic process | 1.62E-05 | 2.23E-03 | | 2/26 7.6% | 7/28950 0.0% | Q99KK2 Q8K297 |
| 44281 | small molecule metabolic process | 1.61E-05 | 2.23E-03 | | 8/26 30.7% | 1350/28950 4.6% | P10649 P04117 Q9Z2I9 O35386 Q99KK2 O09172 Q8VDN2 P57746 |
| 19752 | carboxylic acid metabolic process | 8.67E-05 | 5.79E-03 | | 5/26 19.2% | 518/28950 1.7% | P04117 Q9Z2I9 O35386 Q99KK2 O09172 |
| 43436 | oxoacid metabolic process | 8.67E-05 | 5.79E-03 | | 5/26 19.2% | 518/28950 1.7% | P04117 Q9Z2I9 O35386 Q99KK2 O09172 |
| 6082 | organic acid metabolic process | 8.75E-05 | 5.79E-03 | | 5/26 19.2% | 519/28950 1.7% | P04117 Q9Z2I9 O35386 Q99KK2 O09172 |
| 42180 | cellular ketone metabolic process | 9.83E-05 | 5.79E-03 | | 5/26 19.2% | 532/28950 1.8% | P04117 Q9Z2I9 O35386 Q99KK2 O09172 |
| 33692 | cellular polysaccharide biosynthetcic metabolic process | 1.31E-04 | 6.77E-03 | | 2/26 7.6% | 19/28950 0.0% | Q99KK2 Q8K297 |
|  | MOLECULAR FUNCTION | | | | | | |
| GO-ID | Description | p-val | corr p val | cluster frequency | | total frequency | genes |
| 3756 | protein disulfide isomerase activity | 1.75E-05 | 1.08E-03 | 2/27 7.4% | | 7/28958 0.0% | P27773 Q922R8 |
| 16864 | intracellular oxidoreductase actvity, transposing S-S bonds | 1.75E-05 | 1.08E-03 | 2/27 7.4% | | 7/28958 0.0% | P27773 Q922R8 |
| 16862 | intracellular oxidoreductase actvity, interconverting keto-enol groups | 2.34E-05 | 1.08E-03 | 2/27 7.4% | | 8/28958 0.0% | P27773 Q922R8 |
|  | CELLULAR COMPONENT | | | | | | |
| GO-ID | Description | p-val | corr p val | cluster frequency | | total frequency | genes |
| 44444 | cytoplasmic part | 6.50E-07 | 4.68E-05 | 15/27 55.5% | | 4194/29344 14.2% | P09528 P27773 P04117 Q6RHW0 Q8VDN2 Q9JKR6 P62245 Q9Z2I9 O35386 O09172 Q9D8S9 O89103 Q8K297 P47758 Q922R8 |
| 5737 | cytoplasm | 1.07E-05 | 3.84E-04 | 17/27 62.9% | | 6766/29344 23.0% | P09528 P27773 P10649 P04117 Q6RHW0 Q8VDN2 Q9JKR6 P62245 Q7TSC1 Q9Z2I9 O35386 O09172 Q9D8S9 O89103 Q8K297 P47758 Q922R8 |
| 5622 | intracellular | 1.77E-05 | 4.25E-04 | 20/27 74.0% | | 9784/29344 33.3% | P09528 P27773 P10649 P04117 Q6RHW0 Q8R5A6 Q8VDN2 P70206 Q9JKR6 P62245 Q7TSC1 Q9Z2I9 O35386 Q99KK2 O09172 Q9D8S9 O89103 Q8K297 P47758 Q922R8 |
| 44464 | cell part | 8.83E-05 | 1.27E-03 | 23/27 85.1% | | 14257/29344 48.5% | P09528 P27773 P10649 P04117 Q6RHW0 Q8R5A6 Q8VDN2 P70206 P57746 Q9JKR6 P62245 Q7TSC1 P40240 Q9Z2I9 O35386 Q99KK2 O09172 Q9D8S9 O89103 Q8K297 P47758 Q922R8 Q8BM55 |
| 5623 | cell | 8.85E-05 | 1.27E-03 | 23/27 85.1% | | 14258/29344 48.5% | P09528 P27773 P10649 P04117 Q6RHW0 Q8R5A6 Q8VDN2 P70206 P57746 Q9JKR6 P62245 Q7TSC1 P40240 Q9Z2I9 O35386 Q99KK2 O09172 Q9D8S9 O89103 Q8K297 P47758 Q922R8 Q8BM55 |
| 4424 | intracellular part | 2.89E-04 | 3.46E-03 | 18/27 66.6% | | 9548/29344 32.5% | P09528 P27773 P10649 P04117 Q6RHW0 Q8VDN2 Q9JKR6 P62245 Q7TSC1 Q9Z2I9 O35386 Q99KK2 O09172 Q9D8S9 O89103 Q8K297 P47758 Q922R8 |
| 43231 | intracellular membrane bounded organelle | 4.69E-04 | 4.02E-03 | 15/27 55.5% | | 7111/29344 24.2% | P09528 P27773 P04117 Q8VDN2 Q9JKR6 P62245 Q7TSC1 Q9Z2I9 O35386 Q99KK2 Q9D8S9 O89103 Q8K297 P47758 Q922R8 |
| 43227 | membrane bounded organelle | 4.78E-04 | 4.02E-03 | 15/27 55.5% | | 7123/29344 24.2% | P09528 P27773 P04117 Q8VDN2 Q9JKR6 P62245 Q7TSC1 Q9Z2I9 O35386 Q99KK2 Q9D8S9 O89103 Q8K297 P47758 Q922R8 |
| 43229 | intracellular organelle | 5.38E-04 | 4.02E-03 | 16/27 59.2% | | 8082/29344 27.5% | P09528 P27773 P04117 Q6RHW0 Q8VDN2 Q9JKR6 P62245 Q7TSC1 Q9Z2I9 O35386 Q99KK2 Q9D8S9 O89103 Q8K297 P47758 Q922R8 |
| 43226 | organelle | 5.58E-04 | 4.02E-03 | 16/27 59.2% | | 8107/29344 27.6% | P09528 P27773 P04117 Q6RHW0 Q8VDN2 Q9JKR6 P62245 Q7TSC1 Q9Z2I9 O35386 Q99KK2 Q9D8S9 O89103 Q8K297 P47758 Q922R8 |
| 5783 | endoplasmic reticulum | 9.68E-04 | 6.33E-03 | 5/27 18.5% | | 852/29344 2.9% | P27773 Q9JKR6 Q8K297 P47758 Q922R8 |

**Table S3 Down-regulated GO categories in infected macrophages with *L. donovani* promastigotes**

|  | BIOLOGICAL PROCESS | | | | | |
| --- | --- | --- | --- | --- | --- | --- |
| GO-ID | Description | p val | Corr p val | Cluster freq | Total freq | Genes |
| 7033 | vacuole organisation | 3.10E-10 | 1.40E-07 | 5/36 13.8% | 30/28959 0.1% | P24638 P18242 O88531 P29416 O89023 |
| 7040 | lysosome organisation | 4.73E-09 | 1.07E-06 | 4/36 11.1% | 17/28959 0.0% | P24638 O88531 P29416 O89023 |
| 9056 | catabolic process | 5.87E-07 | 7.09E-05 | 9/36 25.0% | 830/28959 2.8% | P23780 P56542 P18242 P14901 O88531 Q9DBU0 Q3TCN2 P29416 O89023 |
| 8152 | metabolic process | 6.29E-07 | 7.09E-05 | 22/36 61.1% | 6472/28959 22.3% | P10605 Q9WV54 P23780 Q6ZWV3 Q91XI1 P56542 Q69Z38 Q9QZD9 P97864 Q62383 P18242 P97821 P14901 P12265 Q80TM9 P24638 O55201 O88531 Q9DBU0 Q3TCN2 P29416 O89023 |
| 44238 | primary metabolic process | 7.33E-06 | 6.61E-04 | 19/36 52.7% | 5588/28959 19.2% | P10605 Q9WV54 P23780 Q6ZWV3 Q91XI1 P56542 Q69Z38 Q9QZD9 P97864 Q62383 P18242 P97821 P12265 Q80TM9 O55201 O88531 Q3TCN2 P29416 O89023 |
| 44248 | cellular catabolic process | 1.13E-05 | 8.51E-04 | 7/36 19.4% | 635/28959 2.1% | P56542 P18242 P14901 O88531 Q9DBU0 P29416 O89023 |
| 6996 | organelle organisation | 3.02E-05 | 1.95E-03 | 8/36 22.2% | 1026/28959 3.5% | Q80TM9 P24638 P97864 Q6P5D8 P18242 O88531 P29416 O89023 |
| 65007 | biological regulation | 6.87E-05 | 3.88E-03 | 20/36 55.5% | 7126/28959 24.6% | P10605 Q61830 Q6R5N8 P56542 Q8VDV3 P97864 Q91YI4 Q62383 Q6P5D8 Q61151 P14901 Q9QUR7 Q6PGG2 Q80TM9 O55201 Q61193 Q8CIM5 O88531 P29416 O89023 |
| 8344 | adult locomotary behavior | 8.00E-05 | 4.01E-03 | 3/36 8.3% | 67/28959 0.2% | Q91YI4 O88531 P29416 |
| 16043 | cellular component organisation | 1.06E-04 | 4.80E-03 | 10/36 27.7% | 1979/28959 6.8% | Q61830 Q80TM9 P24638 P97864 Q91YI4 Q6P5D8 P18242 O88531 P29416 O89023 |
| 9987 | cellular process | 1.67E-04 | 6.87E-03 | 22/36 61.1% | 8929/28959 30.8% | Q61830 P23780 Q6ZWV3 Q91XI1 P56542 Q69Z38 Q9QZD9 P97864 Q91YI4 Q62383 Q6P5D8 P18242 P14901 Q80TM9 P24638 O55201 O70309 O88531 Q9QXH4 Q9DBU0 P29416 O89023 |
| 30534 | adult behaviour | 2.55E-04 | 9.58E-03 | 3/36 8.3% | 99/28959 0.3% | Q91YI4 O88531 P29416 |
|  | MOLECULAR FUNCTION | | | | | |
| Go-ID | Description | p val | Corr p val | Cluster freq | Total freq | Genes |
| 16787 | hydrolase activity | 9.02E-10 | 1.25E-07 | 16/36 44.4% | 2086/28967 7.2% | P10605 Q9WV54 P23780 P56542 P97864 Q62383 P18242 P97821 P14901 P12265 P24638 Q810A7 O88531 Q3TCN2 P29416 O89023 |
| 3824 | catalytic activity | 4.45E-06 | 3.09E-04 | 18/36 50.0% | 4859/28967 16.7% | P10605 Q9WV54 P23780 Q91XI1 P56542 Q69Z38 P97864 Q62383 P18242 P97821 P14901 P12265 P24638 Q810A7 O88531 Q3TCN2 P29416 O89023 |
| 4197 | cysteine type endopeptidase activity | 2.93E-05 | 1.36E-03 | 3/36 8.3% | 48/28967 0.1% | P10605 P97864 P97821 |
| 4175 | endopeptidase activity | 6.66E-05 | 1.87E-03 | 5/36 13.8% | 347/28967 1.1% | P10605 P97864 P18242 P97821 O89023 |
| 3711 | trancsription elongation regulator activity | 6.72E-05 | 1.87E-03 | 2/36 5.5% | 10/28967 0.0% | O55201 Q62383 |
| 5488 | binding | 1.41E-04 | 3.27E-03 | 24/36 66.6% | 10294/28967 35.5% | P10605 Q61830 P23780 Q91XI1 Q6R5N8 Q69Z38 Q9QZD9 Q91YI4 Q62383 Q6P5D8 P18242 P97821 Q61151 P14901 P12265 Q6PGG2 Q80TM9 P24638 Q99N69 Q810A7 O70309 P29416 Q8K221 O89023 |
| 4553 | hydrolase activity, hydrolysing o-glycosyl compounds | 1.74E-04 | 3.45E-03 | 3/36 8.3% | 87/28967 0.3% | P23780 P29416 P12265 |
| 8234 | cysteine-type peptidase activity | 3.20E-04 | 5.56E-03 | 3/36 8.3% | 107/28967 0.3% | P23780 P29416 P12265 |
| 16798 | hydrolase activity acting on glycosyl bonds | 5.53E-04 | 8.40E-03 | 3/36 8.3% | 129/28967 0.4% | P10605 P97864 P97821 |
| 70011 | peptidase activity acting on L aminoacid peptides | 6.04E-04 | 8.40E-03 | 5/36 13.8% | 559/28967 1.9% | P10605 P97864 P18242 P97821 O89023 |
| 8233 | peptidase activity | 7.77E-04 | 9.81E-03 | 5/36 13.8% | 591/28967 2.0% | P10605 P97864 P18242 P97821 O89023 |
|  | CELLULAR COMPONENT | | | | | |
| GO-ID | Description | p val | Corr p val | Cluster freq | Total frequency | Genes |
| 323 | lytic vacuole | 5.10E-23 | 3.14E-21 | 15/40 37.5% | 197/29353 0.6% | P10605 P31996 Q9WV54 P23780 P56542 P18242 P97821 P12265 P17047 P24638 O88531 Q9DBU0 Q3TCN2 P29416 O89023 |
| 5764 | lysosome | 5.10E-23 | 3.14E-21 | 15/40 37.5% | 197/29353 0.6% | P10605 P31996 Q9WV54 P23780 P56542 P18242 P97821 P12265 P17047 P24638 O88531 Q9DBU0 Q3TCN2 P29416 O89023 |
| 5773 | vacuole | 3.20E-22 | 1.31E-20 | 15/40 37.5% | 222/29353 0.7% | P10605 P31996 Q9WV54 P23780 P56542 P18242 P97821 P12265 P17047 P24638 O88531 Q9DBU0 Q3TCN2 P29416 O89023 |
| 44464 | cell part | 3.53E-09 | 8.71E-08 | 37/40 92.5% | 14257/29353 48.5% | Q61830 P31996 P54116 Q9WV54 Q6ZWV3 Q6R5N8 P97864 Q62383 Q6P5D8 P18242 P97821 Q61151 Q6PGG2 Q80TM9 P24638 Q810A7 Q8CIM5 A1L314 O88531 Q9QXH4 O89023 P10605 P23780 P56542 Q9QZD9 Q91YI4 P14901 P12265 P17047 Q99N69 O55201 Q61193 O70309 Q9DBU0 Q3TCN2 P29416 Q8K221 |
| 5623 | cell | 3.54E-09 | 8.71E-08 | 37/40 92.5% | 14258/29353 48.5% | Q61830 P31996 P54116 Q9WV54 Q6ZWV3 Q6R5N8 P97864 Q62383 Q6P5D8 P18242 P97821 Q61151 Q6PGG2 Q80TM9 P24638 Q810A7 Q8CIM5 A1L314 O88531 Q9QXH4 O89023 P10605 P23780 P56542 Q9QZD9 Q91YI4 P14901 P12265 P17047 Q99N69 O55201 Q61193 O70309 Q9DBU0 Q3TCN2 P29416 Q8K221 |
| 267 | cell fraction | 6.68E-05 | 1.37E-07 | 12/40 30.0% | 1021/29353 3.4% | P10605 P17047 P23780 Q80TM9 P24638 P97864 P18242 P14901 O88531 P12265 Q8K221 O89023 |
| 44444 | cytoplasmic part | 1.42E-08 | 2.50E-07 | 21/40 52.5% | 4194/29353 14.2% | P10605 P31996 Q9WV54 P23780 Q6ZWV3 P56542 Q9QZD9 P97864 Q91YI4 P18242 P97821 P14901 P12265 P17047 Q80TM9 P24638 O88531 Q9DBU0 Q3TCN2 P29416 O89023 |
| 5625 | soluble fraction | 1.98E-07 | 3.04E-06 | 7/40 17.5% | 312/29353 1.0% | P10605 P23780 P97864 P18242 P12265 Q8K221 O89023 |
| 5622 | intracellular | 4.65E-07 | 6.36E-06 | 29/40 72.5% | 9784/29353 33.3% | P31996 Q9WV54 Q6ZWV3 P97864 Q62383 Q6P5D8 P18242 P97821 Q61151 Q6PGG2 Q80TM9 P24638 Q810A7 O88531 O89023 P10605 P23780 P56542 Q9QZD9 Q91YI4 P14901 P12265 P17047 Q99N69 O55201 Q61193 Q9DBU0 Q3TCN2 P29416 |
| 5737 | cytoplasm | 5.80E-07 | 7.13E-06 | 24/40 60.0% | 6766/29353 23.0% | P10605 P31996 Q9WV54 P23780 Q6ZWV3 P56542 Q9QZD9 P97864 Q91YI4 P18242 P97821 Q61151 P14901 P12265 P17047 Q80TM9 P24638 Q99N69 Q810A7 O88531 Q9DBU0 Q3TCN2 P29416 O89023 |
| 44424 | intracellular part | 6.17E-06 | 6.89E-06 | 27/40 67.5% | 9548/29353 32.5% | P31996 Q9WV54 Q6ZWV3 P97864 Q62383 Q6P5D8 P18242 P97821 Q61151 Q80TM9 P24638 Q810A7 O88531 O89023 P10605 P23780 P56542 Q9QZD9 Q91YI4 P14901 P12265 P17047 Q99N69 O55201 Q9DBU0 Q3TCN2 P29416 |
| 43231 | intracellular bounded organelle | 2.85E-05 | 2.77E-04 | 22/40 55.0% | 7111/29353 24.2% | P10605 P31996 Q9WV54 P23780 P56542 Q91YI4 Q62383 Q6P5D8 P18242 P97821 P14901 P12265 P17047 Q80TM9 P24638 O55201 Q810A7 O88531 Q9DBU0 Q3TCN2 P29416 O89023 |
| 43227 | membrane bounded organelle | 2.93E-05 | 2,77E-04 | 22/40 55.0% | 7123/29353 24.2% | P10605 P31996 Q9WV54 P23780 P56542 Q91YI4 Q62383 Q6P5D8 P18242 P97821 P14901 P12265 P17047 Q80TM9 P24638 O55201 Q810A7 O88531 Q9DBU0 Q3TCN2 P29416 O89023 |
| 43229 | intracellular organelle | 6.50E-05 | 5.52E-04 | 23/40 57.5% | 8082/29353 27.5% | P10605 P31996 Q9WV54 P23780 Q6ZWV3 P56542 Q91YI4 Q62383 Q6P5D8 P18242 P97821 P14901 P12265 P17047 Q80TM9 P24638 O55201 Q810A7 O88531 Q9DBU0 Q3TCN2 P29416 O89023 |
| 43226 | organelle | 6.86E-05 | 5.52E-04 | 23/40 57.5% | 8107/29353 27.6% | P10605 P31996 Q9WV54 P23780 Q6ZWV3 P56542 Q91YI4 Q62383 Q6P5D8 P18242 P97821 P14901 P12265 P17047 Q80TM9 P24638 O55201 Q810A7 O88531 Q9DBU0 Q3TCN2 P29416 O89023 |
| 5624 | membrane fraction | 7.18E-05 | 5.52E-04 | 7/40 17.5% | 768/29353 2.6% | P17047 Q80TM9 P24638 P97864 P14901 O88531 P12265 |
| 5626 | insoluble fraction | 9.63E-05 | 6.96E-04 | 7/40 17.5% | 805/29353 2.7% | P17047 Q80TM9 P24638 P97864 P14901 O88531 P12265 |
| 8305 | integrin complex | 5.76E-04 | 3.94E-03 | 26/29353 0.0% | 26/29353 0.0% | O70309 Q9QXH4 |
